# Exploration of RNA outside segmented cells in spatial transcriptomics reveals extrasomatic RNA organization

**DOI:** 10.64898/2025.12.07.692889

**Authors:** Sergio Marco Salas, Michael Dammann, Raphael Kfuri Rubens, Francesca Drummer, Lennard Halle, Sören Becker, Fabian J. Theis

## Abstract

Image-based spatial transcriptomics (iST) enables visualization of RNA molecules in their spatial context, yet up to 40% of transcripts remains unassigned to cells and has been largely overlooked. In this study, we systematically analyze unassigned RNAs (uRNAs) across 14 public iST datasets and multiple technologies to characterize their nature and relevance in tissue biology. By assessing potential technical origins, in particular segmentation errors, noise, and diffusion across many tissues in both humans and mice, we find that around one third of uRNAs cannot be attributed to technical artifacts. Those non-technical uRNAs are enriched around cells with complex morphologies such as neurons, glia, and endothelial cells and reflect transcripts localized in cellular protrusions and extrasomatic compartments. Using these signals, we infer protrusion-associated transcript localization and identify cell-cell contacts beyond standard cell-centric segmentation. Our results challenge the assumption that uRNA is purely technical noise and instead highlight its potential biological relevance, particularly in relation to intracellular RNA localization and tissue architecture. To enable their systematic study, we introduce troutpy, a Python package for quantitative uRNA exploration in spatial transcriptomics data.

## Introduction

Spatial transcriptomics (ST) has revolutionized molecular biology by enabling researchers to visualize and quantify RNA molecules within their spatial context ^1,2^. The secutting-edge technologies generate high-resolution maps of gene expression patterns in tissues, revealing cell organization and advancing our understanding of pathological processes and disease mechanisms^3^. Spatial transcriptomics methods can be classified into two main families based on the strategy used to profile spatial gene expression: sequencing-based and imaging-based (iST)^4,5^. Sequencing-based methods (e.g. Visium^6^, Stereo-seq^7^) attach spatial barcodes to transcripts, allowing spatial assignment after sequencing. In contrast, imaging-based methods (e. g., Xenium^8^, MERSCOPE^9^, CosMx^10^) directly visualize individual RNA molecules in situ through high-resolution imaging, allowing precise subcellular localization. Consequently, while sST assigns transcripts to spatial positions via barcodes, iST directly measures the exact positions of individual molecules.

Since iST methods retrieve the location of individual transcripts, they usually rely on cell segmentation to define cell boundaries and assign transcripts accurately. In recent years, multiple methods have been developed for this task, including strategies based on complementary stainings^11,^, transcript location^13,^ and hybrid approaches^15,^. Though no benchmark exists, multiple studies have recently highlighted how introducing transcript information improves segmentation performance. Despite these advances, cell segmentation presents significant challenges, including accurately delineating cell borders in densely packed tissues, distinguishing between overlapping cells^17^, and accounting for variability in cell morphology. Thus, perfect segmentation in 2D sST datasets is unlikely to be achieved, particularly in complex tissues with fine cellular protrusions. These complexities can lead to segmentation errors, impacting downstream analyses of intracellular RNA^18,19^.

During segmentation, a substantial proportion of detected transcripts remain unassigned to any segmented cell^18,20,21^. These molecules, referred to as unassigned RNA (uRNA) throughout this manuscript, are systematically omitted in the analysis of iST data. The uRNA pool may have heterogeneous origins, potentially arising from technical (e.g. missegmented cells, noise, RNA diffusion) or biological sources (e.g. cellular protrusions, projections, RNA in vesicles). Despite being largely overlooked, several studies have noted the presence and potential significance of uRNA but not significantly followed up on their origin and meaning. Technology benchmarking studies suggest that forcing all transcripts into cells can reduce segmentation accuracy^18,20^. Similarly, a recent study^13^ introduced the concept of *fragments*, groups of transcripts with local and spatial coherence that lack an associated nucleus, and detected their presence in human breast cancer samples profiled with Xenium. In the mouse brain, groups of synapse-associated mRNAs have been found outside segmented cells^22^, revealing that uRNA can mark synaptic organization and provide spatial insights that extend beyond traditional cell boundaries21.

These observations challenge the assumption that uRNA is merely technical, instead suggesting it may carry biologically meaningful signals worth further investigation. However, current approaches only capture certain aspects of uRNAs, describing their occurrence indirectly rather than systematically characterizing their features or spatial distribution. Furthermore, it remains unclear whether these signals are technical or biologically meaningful. The refore, a comprehensive characterization is needed to better understand the nature of these transcripts, in particular whether they are purely technical or carry biological meaning. In this study, we therefore systematically investigate uRNA across multiple image-based spatial transcriptomics platforms and tissues. We analyze its spatial distribution, explore potential biological roles, and evaluate associated technical factors. To support broader exploration of uRNA across biological systems, we present troutpy (Extended Data Figure 1A), an open-source package in Python for the analysis and interpretation of unassigned RNA in spatial transcriptomics data.

## Results

### Missegmentation is a major source of unassigned RNA

A comprehensive understanding of unassigned RNA (uRNA) requires examining its occurrence and characteristics across diverse biological contexts and technical platforms. For this, we compiled a collection of 14 public iST datasets^23-26^ encompassing multiple organs, species, and technologies (Figure 1A, Extended Data Table 1). Using default segmentation, 39.1% of transcripts were unassigned on average (Figure 1B), ranging from 4% to 84.2% across tissues and datasets. Segmentation strategy, more than tissue type, drove variability, with nuclei-based methods producing higher u RNA than multimodal approaches. This is exemplified in the mouse brain datasets, where 61.3% of transcripts were unassigned with nuclei-based segmentation versus 35.1 ± 1. 11% with multimodal segmentation (Figure 1B). To evaluate whether more accurate segmentation could reduce uRNA, we applied Proseg^14^, a representative state-of-the-art segmentation strategy, to mouse brain datasets as an exemplar of advanced segmentation approaches.. While this strategy resulted in a lower proportion of uRNA, a substantial fraction of transcripts remained unassigned (mean = 25.5%) (Extended Data Figure 1B). Importantly, the composition of these unassigned transcripts remained largely unchanged (Extended Data Figure 1C-E), showing that their profile is consistent and that they persist even with current state-of-the-art segmentation methods.

**Figure 1.**
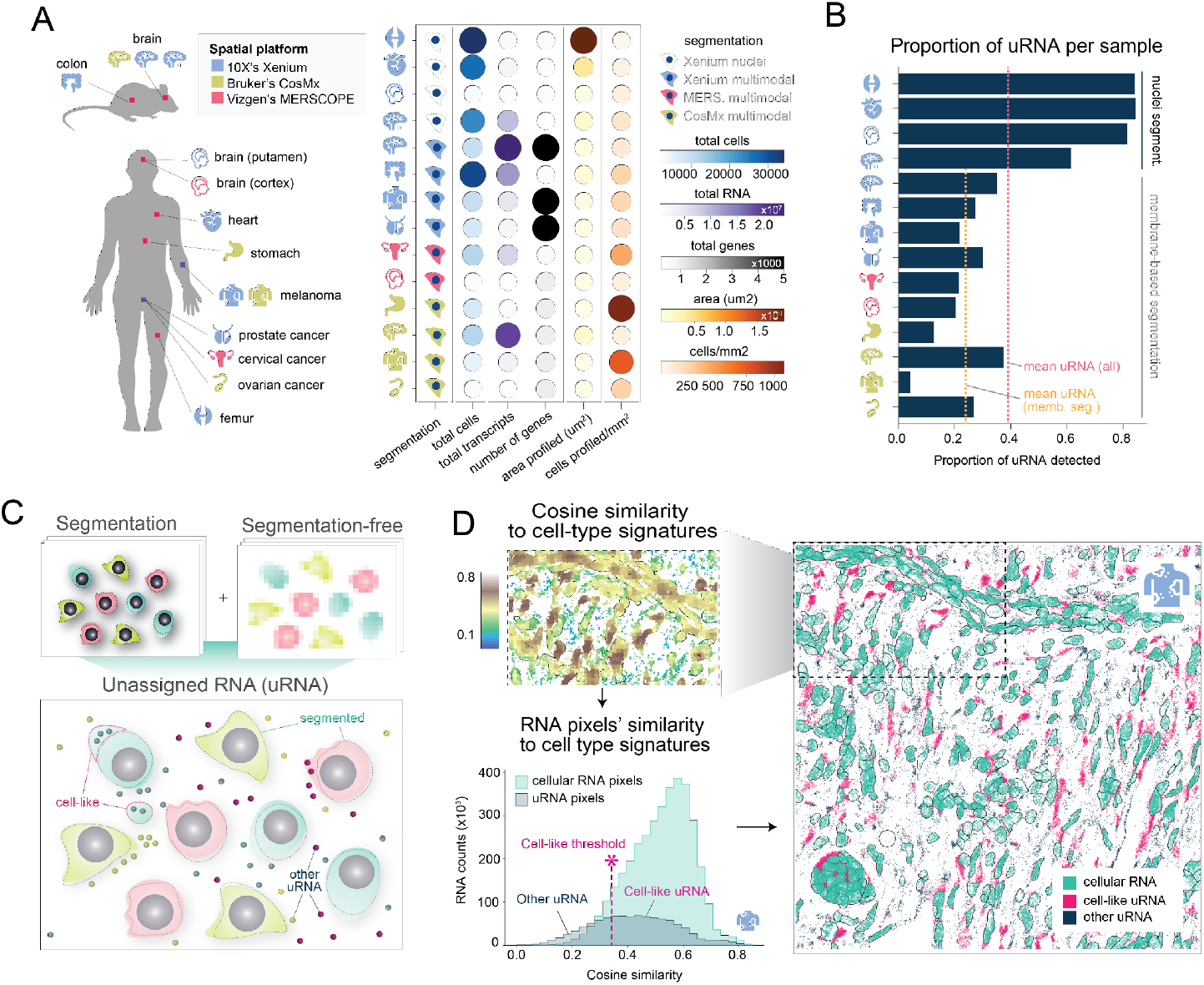
Quantification of unassigned RNA reveals tissue variability. **(A)** Schematic representation of the dataset collection used during the study (left) and their main characteristics (right). **(B)** Bar plot representing the proportion of unassigned RNA within each sample. **(C)** Schema of the approach followed to identify unassigned RNA by combining segmentation and segmentation-free strategies. **(D)** Workflow used to identify mis-segmented regions using the combination of segmentation and segmentation-free strategies, exemplified in a Xenium melanoma region of interest.

Although state-of-the-art segmentation strategies can improve transcript assignment, their effectiveness remains uncertain due to lack of systematic benchmarking. As a consequence, even when applying them, some cellular structures might remain unsegmented, contributing to the uRNA pool. To detect these, we developed a customized strategy combining segmentation’s output and Sainsc^27^, a segmentation-free approach. Briefly, the tissue is divided into equally sized pixels, and each pixel’s transcriptomic profile is compared to predefined cell type signatures. Pixels outside segmented cells that show similarity comparable to pixels within cells are classified as cell-like unassigned regions, capturing structures missed by segmentation (Figure 1C-D, Methods). This approach was robust to hyperparameter tuning, producing consistent uRNA signatures (Extended Data Figure 2 A-C). By applying this strategy across datasets, we found that on average 62.4% of uRNA was located in cell-like unassigned regions (Figure 2E), indicating that missegmentation is a major contributor. Substantial tissue-specific differences were observed, showing that the impact of missegmentation depends on tissue and cell type. By characterizing for example these cell-like uRNA regions in a Xenium mouse brain dataset (Extended Data Figure 2D-H), we found they are unevenly distributed across cell types, with most segments located adjacent to segmented cells and some extending beyond segmentation boundaries, compatible with biological structures missed by segmentation.

**Figure 2.**
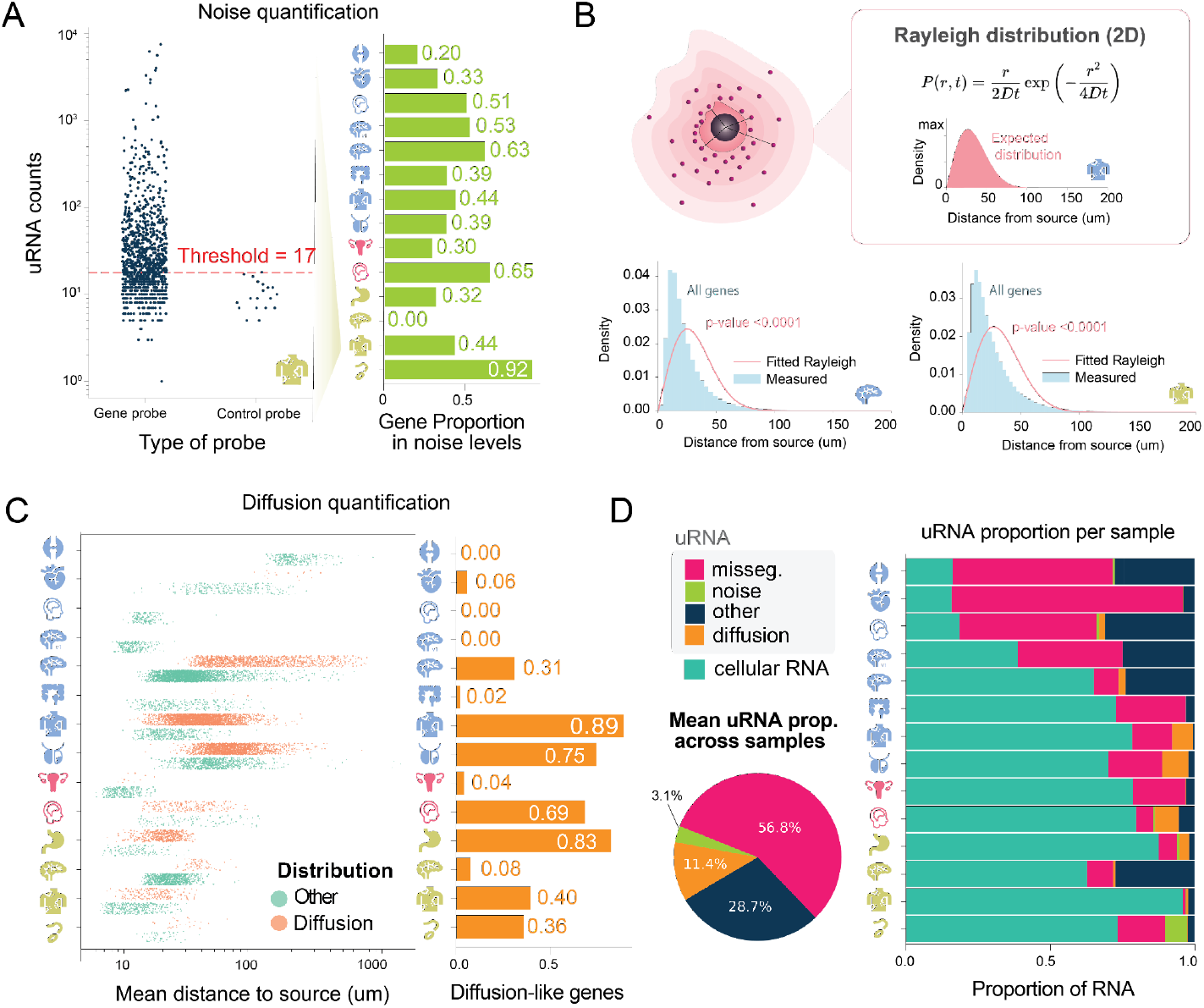
Quantitative evaluation of uRNA origins reveals consistent non-technical uRNA. **(A)** Quantification of the number of genes contributing to the uRNA pool with counts below the noise levels, per sample (left). For the CosMx melanoma sample, the number of counts in the uRNA pool for both gene probes and control probes, as well as the selected noise threshold, is presented (right). **(B)** Schematic representation of the distribution of RNA expected from 2D diffusion, modeled by the rayleigh distribution (top). Examples of the fitted rayleigh distribution and the measured distribution in regards to the distance from each transcript to their closest source is shown for two samples: mouse brain profiled with Xenium Prime (bottom, left) and melanoma profiled with CosMx (bottom, right). P-values derived from the Kolmogorov-Smirnov test applied to compare the Rayleigh fitted and the measure distribution are shown for both samples. **(C)** Proportion genes exhibiting a uRNA distribution compatible with diffusion per sample (left) and gene-level assessment for the CosMx melanoma sample (right). This plot is complemented by a strip plot representing the mean distance to source (x-axis) for each gene profiled by sample (y-axis) (left). Genes are colored based on whether their distribution to source is different from the diffusion-compatible Rayleigh distribution (diffusion) or not (other). **(D)** Stacked bar (right) plot representing, for each sample, the proportion of RNA segmented into cells (cellular RNA) and the unassigned proportion (uRNA), grouped by their predicted origin, including missegmentation (misseg.), noise and diffusion. All uRNA with an unidentified technical origin are presented as “other”. A pie plot representing the mean uRNA proportions belonging to each category across samples is also included (bottom, left)

### Quantifying the impact of noise and diffusion on unassigned RNA

A notable percentage of uRNA (mean=43.2 ± 22.8%) (Figure 2D) can not be explained by missegmentation and thus, may have a different origin. Multiple technical uRNA sources are proposed as a source of uRNA^28,29^, primarily technical noise and RNA diffusion. To assess technical noise, we used negative control probes, available in some iST platforms. These probes vary in design, but are generally intended to produce no signal, either because they do not hybridize to any endogenous RNA or because their decoding scheme does not match any measured gene. Therefore, detected signals reflect background noise, defining the experiment’s noise threshold (Extended Data Figure 3A, Methods). For platforms lacking such probes (e.g., MERSCOPE), we developed a custom approach to estimate noise levels (see Methods). Genes with uRNA detection levels below this threshold are likely explained by technical noise (see Methods). Across datasets, on average, 42% of profiled genes per sample exhibited an uRNA abundance below their noise threshold, with a wide tissue variability (Figure 2A).

Next, we evaluated the contribution of RNA diffusion to the uRNA pool. In this context, we define RNA diffusion as the passive spread of transcripts from cellular origin to nearby locations where they are detected, driven mainly by random motion. Diffusion is an extensively reported phenomenon in several sST platforms^6,30^ and multiple methods have been developed to mitigate its effect in downstream analysis^31,32^. However, this phenomenon is less characterized in iST.

While diffusion in tissue sections is inherently three-dimensional, the geometry makes it complex: sections are only 5-10 μm thick, with one open surface allowing outward diffusion and one closed surface against the glass slide that likely restricts movement. This pseudo-3D setup complicates modeling, though in high-resolution platforms we often observed a mild accumulation of unsegmented transcripts near the slide surface, consistent with diffusion-limited trapping ( Extended Data Figure 3B). Given these challenges, and for the sake of tractability, we modeled diffusion as a two-dimensional process, focusing on lateral RNA displacement from source cells. Assuming uRNA originates from nearby expressing cells, we assigned each unsegmented transcript to the nearest cell expressing the same gene and measured their distance to assess consistency with passive diffusion (Methods). In this 2D system, passive diffusion from a point source would be expected to produce a Rayleigh distribution^33^ of uRNA-source distances (Figure 2B, Methods). We used the Kolmogorov-Smirnov test to compare uRNA-source distances with the expected Rayleigh distribution. Simulations confirmed this approach accurately captures diffusion-derived patterns (Extended Data Figure 3C-D, Methods). Across datasets, we found that the empirical distributions significantly deviated from the Rayleigh expectation (Figure 2B), suggesting that passive diffusion alone does not fully account for the global uRNA patterns observed. Nonetheless, the contribution of passive diffusion varies by gene. To investigate this, we compared the distance distribution of uRNA for each individual gene to the Rayleigh model. On average, 31.1% of all genes showed distributions consistent with Rayleigh expectations across datasets (Figure 2C, Extended Data Figure 3D-E). Importantly, the mean distances between uRNAs and their predicted source cells per gene varied widely.

Overall, our analysis shows that the majority (56.8%) of uRNA is linked to missegmentation, while technical noise accounts for only 3.1% and diffusion-compatible patterns represent only 11.4% of the signals. Notably, the remaining 28.7% of uRNA could not be explained by any of the proposed technical origins. This uRNA is particularly abundant in tissues with complex cell morphology like the brain or bone.

### Gene-Specific uRNA distributions reveal distinct origins and behaviors

To understand the uRNAs not linked to technical artifacts, we examined their characteristics. We found substantial variability in uRNA abundance and proportion (the fraction of transcripts classified as uRNA per gene), with some genes especially enriched in the uRNA pool (Figure 3A). For example, several keratin genes (KRT1, KRT10, KRT80, KRT17), expressed in stratified squamous epithelium, showed a high uRNA proportion (> 30%) in the CosMx melanoma dataset. (Figure 3A-B, Extended Data Figure 3E). In the mouse colon sample, Gfap, expressed in enteric glial cells^34^, was the gene with the highest uRNA proportion (Figure 3 C-D). In both cases, these genes are expressed by cell types presenting irregular morphologies. Furthermore, the distribution of these unassigned transcripts lies in proximity to cells expressing these genes (Figure 3 B,D), positioning missegmentation as a likely source. However, these transcripts are not identified as cell-like when assessing missegmentation, indicating that their local molecular composition differs from that of the corresponding cell bodies. This pattern may result from active mRNA transport, where specific RNAs are directed to particular subcellular structures, creating spatial RNA signatures that differ from cell expression profiles. Consistent with this idea, we observed variation in uRNA abundance among genes expressed by the same cell type. For enteric glial cell markers, Sox2 and Plp1 had about 10% of transcripts classified as unassigned, while Gfap showed a much higher proportion at 23.3%. This aligns with previous knowledge, as Sox2, a nuclear transcription factor, is expected to remain in the nucleus, whereas Gfap is known to localize to glial processes even at the RNA level, contributing to cytoskeletal structure^35^.

**Figure 3.**
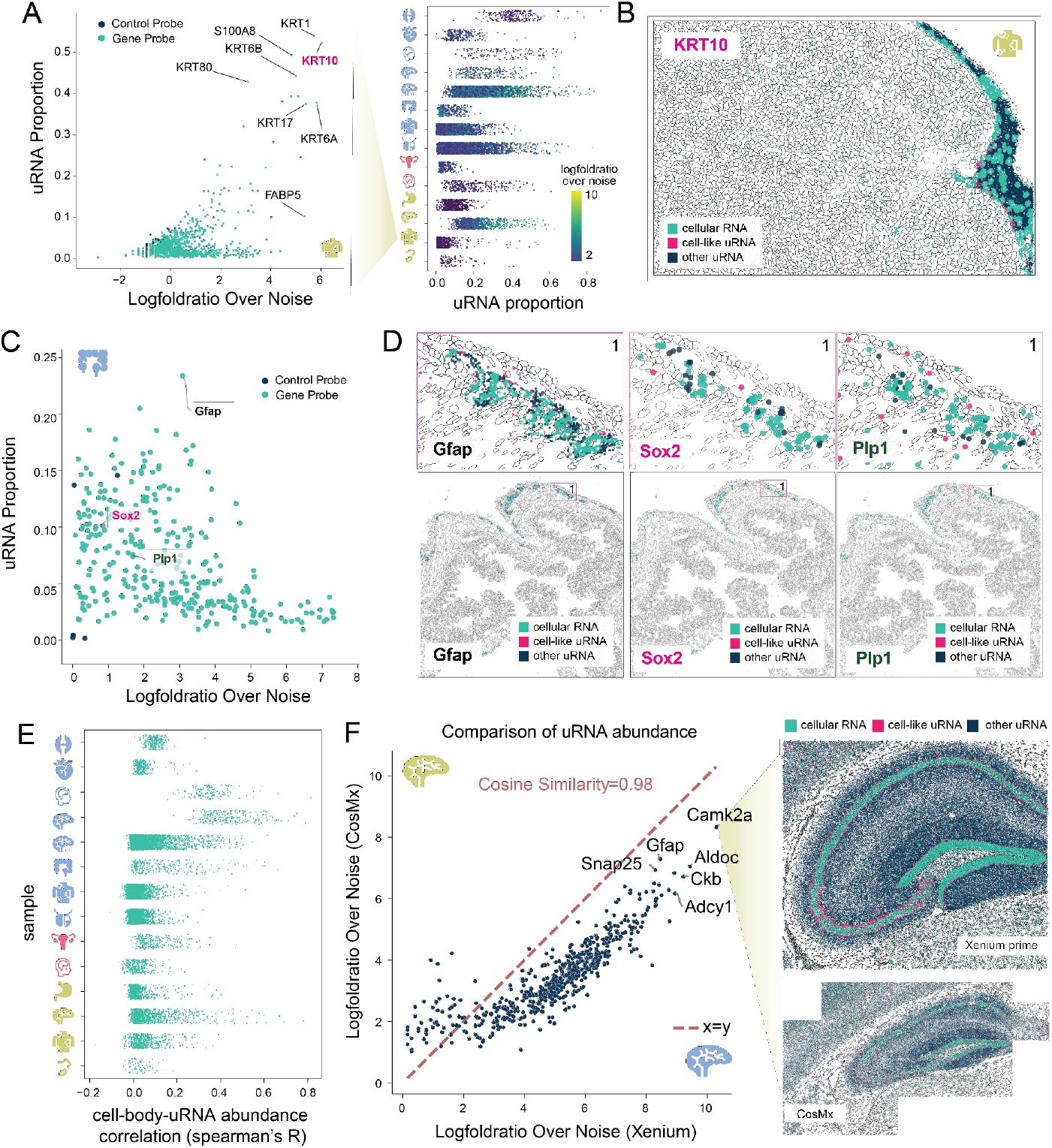
Exploration of uRNA characteristics highlight gene-specific uRNA features. **(A)** Strip plot showing the uRNA proportion (x-axis) for each gene (across all samples, y-axis). Points are colored by the uRNA abundance of each gene, expressed as the log-fold ratio over a noise threshold (color scale, right). A corresponding scatter plot is shown for the CosMx melanoma sample, with uRNA abundance (log-fold ratio over noise) on the x-axis and uRNA proportion on the y-axis. The name of selected genes presenting a high uRNA proportion is highlighted. **(B)** Spatial map of the CosMx melanoma sample. Transcripts assigned to KRT10, one of the genes with the highest uRNA abundance and proportion (see panel A), are displayed and colored by RNA status: cellular RNA, or uRNA. uRNA is further subdivided based on spatial context into cell-like uRNA (found in cell-like regions) and other uRNA. Cell boundaries from the default CosMx segmentation are overlaid in the background. **(C)** Scatter plot of the genes profiled in the mouse colon using Xenium representing their uRNA abundance (log-fold ratio over noise) on the x-axis and uRNA proportion on the y-axis. Genes expressed in glial enteric cells and visualized in panel D are highlighted. **(D)** Spatial maps of the Xenium mouse colon sample showing both the fully analyzed area (bottom) and a zoomed-in region of interest labeled as 1 (top). The maps display transcripts for three enteric glial cell markers: Gfap (left), Sox2 (middle), and Plp1 (right). Transcripts are colored by RNA status: cellular RNA, or uRNA. uRNA is further categorized based on spatial context into cell-like uRNA (located in regions resembling cells) and other uRNA. Cell boundaries from Xenium’s multimodal segmentation are shown in the background for reference. **(E)** Strip plot showing the cell-body-uRNA expression correlation for each gene, calculated as the Spearman correlation (R) between expression inside segmented cells and in the surrounding unsegmented space (x-axis), across samples (y-axis). **(F)** Scatter plot comparing gene-level uRNA abundance between biologically comparable hippocampal regions in the Xenium (x-axis) and CosMx (y-axis) mouse brain samples (left). uRNA abundance is represented as the log-fold ratio over background noise for each gene. A diagonal reference line (x = y) is included to aid visual comparison. The names of the top five genes with the highest uRNA abundance in both samples are labeled. Spatial maps of the gene Camk2a, the most abundant uRNA gene in both datasets, are shown for the Xenium sample (top right) and the CosMx sample (bottom right). Transcripts are colored by RNA status: cellular RNA or uRNA. uRNA is further subdivided into cell-like uRNA (found in regions resembling cells) and other uRNA. Cell boundaries derived from each platform’s segmentation are shown in the background.

We next explored whether, for each gene, their spatial patterns in segmented cells matched their uRNA counterparts. We identified consistently low correlations, indicating that uRNAs often occupy distinct spatial regions (Figure 3E). A clear example is Ppp1r16b, which presented a high uRNA spatial variability in the mouse brain (Moran’s I = 0.33), forming a distinct spatial pattern compared to its cell body expression (Spearman’s R = 0.15). In this case, while most of its uRNA is localized in the dentate gyrus molecular layer (DG-mo), expressing cells reside in the granule layer (DG-sg) (Extended Data Figure 3F), in agreement with its reported enrichment in neurons’ dendrites^36^.

If the origin of these transcripts is biological, their abundance in the uRNA pool should be consistent across platforms. To test this, we compared the uRNA composition of the mouse hippocampus using biologically equivalent samples profiled with two different iST platforms (Xenium and CosMx). Importantly, we found strong cross-platform correlations in u RNA abundance, spatial autocorrelation, and uRNA proportion for individual genes, indicating that the same genes were consistently detected, similarly organized in space, and present in comparable proportions relative to their total expression (Figure 3F, Extended Data Figure 3G). Notably, genes associated with synaptic plasticity and known to localize to cellular protrusions^37^, (i.e. *Camk2a, Ckb, Snap25)*, were among the most enriched genes in both uRNA pools. This cross-platform consistency indicates that non-technical uRNA signals are robust and likely reflect genuine biology rather than artifacts.

### Spatial patterns of uRNA reflect extrasomatic architecture

Certain genes display distinctive spatial characteristics in their uRNA portion. We hypothesized that these patterns might reflect coordinated expression programs. To explore this, we focused on a Xenium prime mouse brain dataset (Extended Data Table 1), dividing the tissue into uniformly spaced 10 × 10 μm square bins and grouping uRNA within each bin. This naïve approach avoided assumptions about cellular or anatomical structure, unlike other strategies^13,22^. uRNA clusters, identified based on molecular similarity, aligned with known brain anatomical structures and did not generally correspond with the expression or location of any of the cell types (Figure 4A-B).

**Figure 4.**
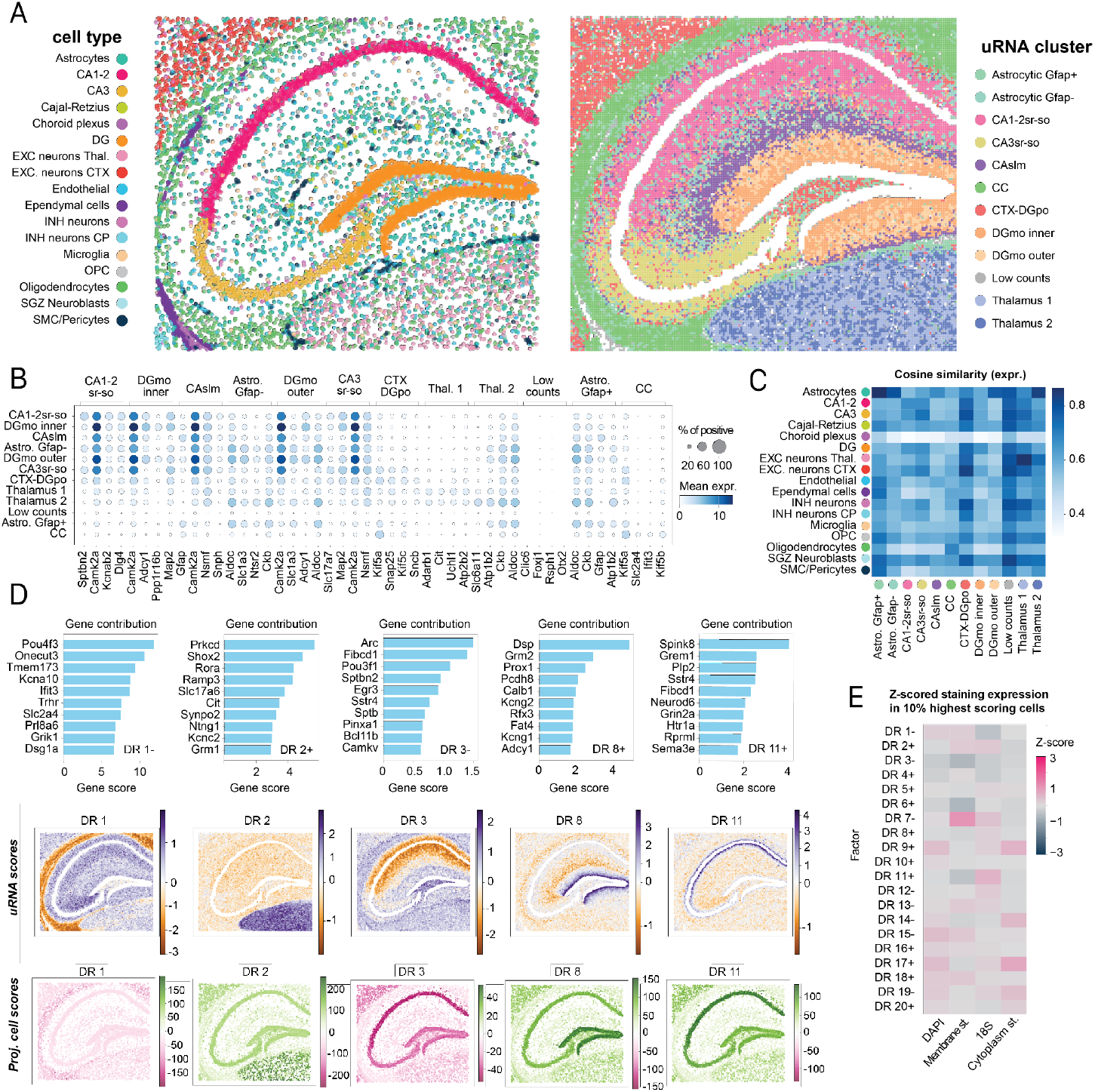
uRNA patterns in the mouse brain reflect an organized extrasomatic architecture. **(A)** Spatial map of the cell types (left) and uRNA clusters (right) identified in the mouse brain Xenium sample. **(B)** Dot plot visualizing, for each uRNA cluster annotated, the expression of its top 3 differentially expressed genes, across uRNA clusters. Dot size represents the fraction of positive bins per gene, whereas the color indicates the mean expression of the gene in the uRNA cluster. **(C)** Heat map representing the cosine similarity between the mean expression signatures of the annotated cell types in panel A (y-axis) and the uRNA clusters (x-axis) **(D)** Gene contribution map (top), spatial map of uRNA bins colored by DR score (mid) and projected DR scores in segmented cells (bottom) of a set of selected DRs, each of them represented in one column. **(E)** Heatmap showing the Z-score of staining intensities (x-axis) across complementary Xenium stainings, calculated within the top 10% most positive bins for each DR (y-axis), either in the positive (+) or negative (−) direction. Membrane staining (membrane st.) represents the cocktail used in the Xenium multimodal staining composed by ATP1A1,CD45 and E-cadherin, while cytoplasmic staining (cytoplasm st.) represents the cocktail composed by AlphaSMA and Vimentin.

To test whether these signatures simply reflected admixture from nearby cells, we applied non-negative matrix factorization (NMF) to identify underlying biological factors driving the uRNA heterogeneity (Extended Data Figure 5). Several of these factors captured interpretable and spatially distinct biological patterns that extended beyond simple admixture artifacts, with most showing limited alignment to specific cell types and marked enrichment in subcellular compartments. While NMF is a powerful approach, its linear nature limits the ability to disentangle overlapping sources of variability, motivating the use of non-linear approaches. Therefore, we additionally applied DRVI^38^ to disentangle the latent dimensions, referred as DR, driving the uRNA local variability, resulting in an unbiased identification of gene programs driving the uRNA spatial variation (Figure 4 D-E, Extended Data Figure 4). Our analysis revealed a total 20 DRs, some of them corresponding to specific cell types (e.g. DR 10: microglia, DR16: oligodendrocyte precursor cells), consistent with missegmentation. Notably, DR3 and DR11, both linked to CA1-CA2 pyramidal neurons, showed distinct spatial patterns. DR3, driven by Arc^39^ and Fibcd1, spanned the CA1-CA2 field, reflecting dendritic localization, while DR11, marked by Spink8 and Grem1, localized near the pyramidal layer. Finally, some DRs (e.g. DR1: corpus callosum or DR4: DGmo) were region-specific. Complementing this, we found that uRNA variability partially correlated with pixel intensities of the morphological stainings (Figure 4E). For instance, the bins with highest scores for cell type-specific DRs, such as DR11 (CA1-CA2), were often located in areas with higher pixel intensity for certain stainings (i.e. 18S staining). In contrast, region-specific DRs, such as DR1 and DR4, were identified in areas with lower pixel intensities across all stainings. Together, our analysis supports the idea that uRNA can capture biological variability at different scales, spanning from cellular structures to tissue domains.

To demonstrate the relevance of our analysis beyond neural tissue, we applied it to a Xenium femur dataset, identifying uRNA genes and signatures with distinct spatial distributions and molecular profiles that were not captured by segmented cells (Extended Data Figure 6). Examples of this are the uRNA-exclusive OGN and AQP3 patterns identified in the bone periphery and the identification of multiple NMF-derived factors corresponding to uRNA-exclusive gene expression signatures, scoring high in bins located within the bone, where no cells were segmented (Extended Data Figure 6). Our results highlight that structured uRNA organization can occur across tissues, likely related to diverse biological processes.

### Unassigned RNA provides insights into intracellular RNA localization patterns and cellular architecture

As shown earlier, non-technical uRNA provides information complementary to segmented cells. Assuming it originates from cells in the tissue, we wondered if it would be possible to predict the contribution of each cell to the uRNA pool, based on its expression. To achieve this, we developed two complementary metrics. First, we implemented a uRNA contribution score, which estimates, for each gene, the origin of its unassigned transcripts as a weighted sum across all cells expressing the gene. By aggregating across genes, this score reflects the number of uRNAs attributable to each cell based on its expression (see Methods). We exemplified the use of this metric in the Xenium 5K mouse brain dataset (Figure 5A, Extended Data Figure 7A), where this metric highlighted several neuronal subtypes as major uRNA contributors. As expected, when aggregating scores by cell type, the most abundant populations accounted for most of the uRNA signal (Extended Data Figure 7D). To account for differences in total transcript counts, we also computed a normalized uRNA contribution score, representing the expected number of uRNA per intracellular transcript. We can interpret the score as a segmentation quality metric since it reflects, for a given cell, which proportion of its transcripts were assigned to cells. In the mouse brain example, astrocytes and endothelial cells emerged as top contributors (Extended Data Figure 7A-B).

**Figure 5.**
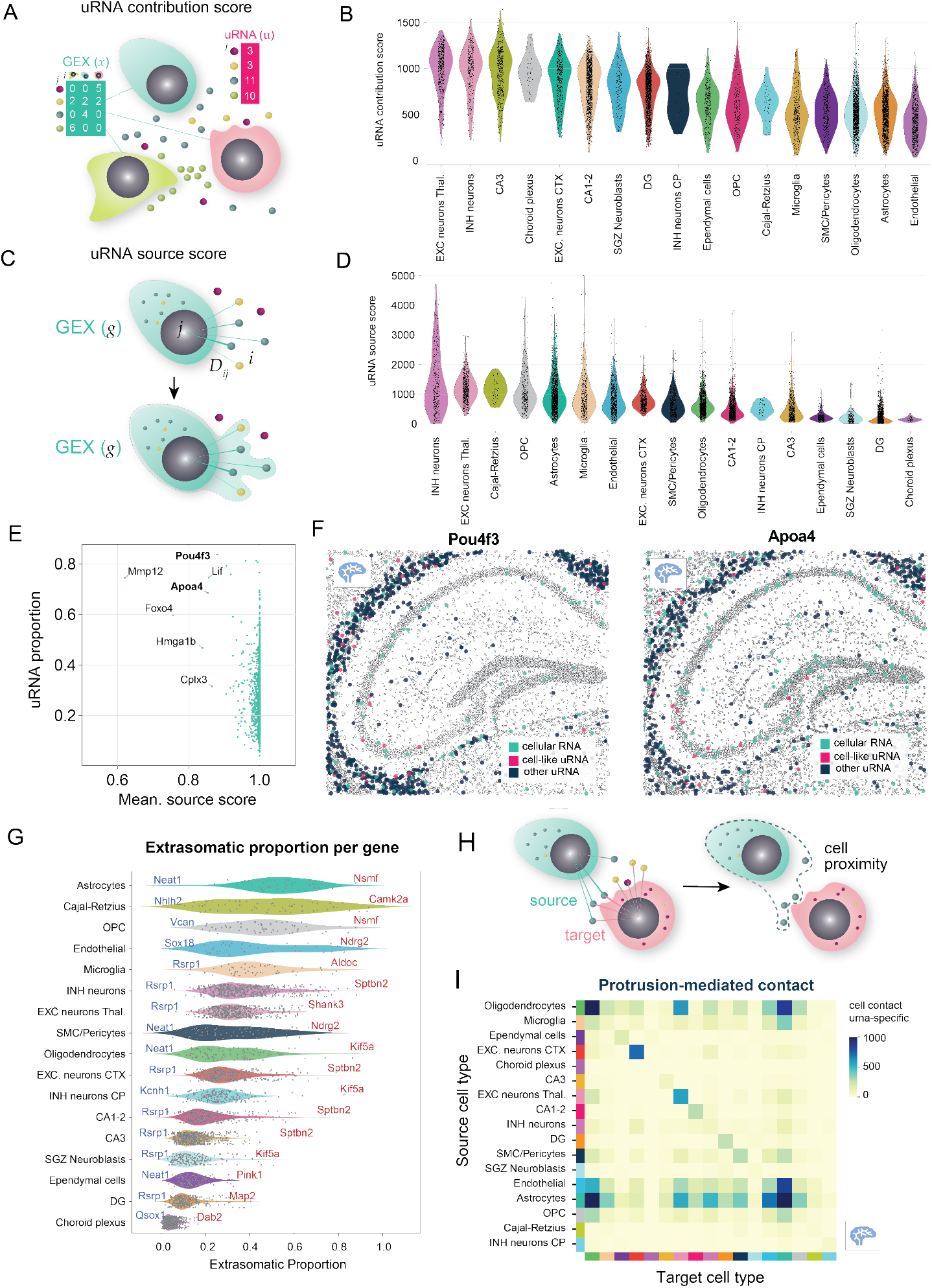
Cell-centric uRNA metrics provide insights into the mouse brain’s intracellular RNA localization patterns and cellular architecture. **(A)** Schema representing the rationale and formula behind the uRNA contribution score. **(B)** Violin plot representing the distribution of the scores by cell type in the Xenium mouse brain dataset (right). A strip plot with the score of each cell, grouped by cell type is overlaid on the violin plot. **(C)** Schematic representation of the rationale and formula behind the uRNA source score. **(D)** Violin plot representing the distribution of the scores by cell type in the Xenium mouse brain dataset. A strip plot with the score of each cell, grouped by cell type is overlaid on the violin plot. **(E)** Scatter plot representing the uRNA proportion (y-axis) and the mean source score for the sum of all cell types (y-axis) for genes that presented an uRNA abundance (log fold ratio over noise threshold of 4). The names of genes with a mean source score below 0.9 are shown. The names of genes shown in F are presented in bold. **(F)** Spatial maps of the Xenium mouse brain 5k sample displaying transcripts for two genes highlighted in panel E: Pou4f3 (left), and Apoa4 (right). Transcripts are colored by RNA status: cellular RNA, or uRNA. uRNA is further categorized based on spatial context into cell-like uRNA (located in regions resembling cells) and other uRNA. Cell boundaries from Xenium’s multimodal segmentation are shown in the background for reference. **(G)** Violin plot showing the distribution of computed extrasomatic proportions for all genes in each cell type. Overlaid is a scatter plot where each point represents the extrasomatic proportion of a specific gene-cell type pair, as computed using troutpy. For each cell type, the names of the genes with the highest and lowest extrasomatic proportions are labeled **(H)** Schematic representation of the rationale behind the use of uRNA to quantify protrusion-mediated cell-cell contact. **(I)** Heatmap showing the number of cell-cell contacts identified using unassigned transcripts.Source cell types (presumed uRNA producers) are shown on the y-axis, and target cell types (closest cells to each uRNA molecule) are on the x-axis.

While this metric can be relevant, it does not consider spatial information, critical to understand the cellular origin of each uRNA. To address this, we developed a second metric, named uRNA source score. Essentially, for each uRNA, a probability of each cell being its source is computed based on each cell’s expression and their spatial proximity weighted by distance. Each cell’s uRNA source score will be the sum of the probabilities across uRNAs (Figure 5B,see Methods). As done for the uRNA contribution score, the source score can be normalized by total counts to reveal the relative uRNA source score of each cell in relation to its expression. As it occurred in the previous metric, endothelial cells and astrocytes were the cell types with a highest normalized uRNA source score (Extended Data Figure 7B). Importantly, normalized contribution and source scores differ across cells, since the contribution score relies only on a cell’s expression and the uRNA pool, while the source score also incorporates spatial information to assign uRNAs to their source cells.

Assigning a source assumes the uRNA originates from a segmented cell, which may not hold for all genes. To account for this, the uRNA source score includes a bias term that captures transcripts without a reliable nearby source cell. By aggregating source scores across cell types for each gene in the mouse brain, we identified genes with unexpectedly low source scores, suggesting a lack of an identifiable cell of origin (Figure 5C). Notable examples include *Pou4f3, Apoa4, Mmp12*, and *Foxo4*, which are robustly detected in the extracellular pool, and which show a high uRNA proportion and yet cannot be traced back to any segmented cell. Interestingly, many of these transcripts are spatially enriched in the corpus callosum, a region rich in myelinated axons, which are poorly captured by segmentation. Some, like *Pou4f3*, are known to be expressed in glutamatergic midbrain^40^ neurons located outside the profiled region.

We next hypothesized that the uRNA source score could be used to estimate the abundance of each transcript type within cellular protrusions across cell types (see Methods). This is particularly valuable given the limitations of current technologies in detecting RNA localized to protrusions^41^. Applying this approach to the Xenium 5K mouse brain dataset, we observed substantial variability in the proportion of extrasomatic transcripts across both genes and cell types. Importantly, these findings align with known biology. For example, *Rsrp1*, which encodes a protein involved in nuclear RNA splicing, showed among the lowest extrasomatic enrichment across multiple cell types. In contrast, *Nsmf*, linked to axon guidance^42^, and *Sptbn2*, associated with dendritic localization^43^, were among the most enriched in the extrasomatic space. To validate our strategy, we leveraged the dataset’s nuclear and cytoplasmic segmentations as ground truth for transcript localization. We found strong concordance between our estimates and the true nuclear-cytoplasmic distribution (Extended Data Figure 7E-F), supporting the accuracy of our approach. However, cytoplasmic transcript abundance was consistently underestimated. This limitation arises because distal transcripts are assigned only if the cell also expresses the gene locally. Transcripts in projections without somatic expression may go unassigned, underestimating protrusion-associated abundance.

As shown throughout this study, a substantial portion of the uRNA pool reflects incomplete segmentation, whether from the soma or peripheral extensions. In this context, unassigned transcripts may serve as markers of complex cell morphology and help reveal aspects of tissue architecture not captured by conventional segmentation. One application of this principle is the study of cell-cell contacts. To explore this, we developed a protrusion-based scoring framework to infer cell-cell interactions from uRNA spatial distribution. For each uRNA, we inferred both its source and target cells (see Methods), using the spatial displacement between them as evidence for protrusion-mediated contact. Aggregating these interactions yielded a directional contact matrix between cell types. When applied to the Xenium mouse brain dataset, this framework recapitulated known cell-cell interactions (Figure 5G-H) and uncovered additional protrusion-mediated contacts not detectable by standard cell-centric approaches (Extended Data Figure 7G-H). Notably, astrocyte-mediated interactions were more prominent using this method, consistent with their known morphological complexity and functional roles in contacting multiple cell types.

## Discussion

Spatial transcriptomics technologies have primarily focused on RNA molecules assigned to segmented cells, often discarding a large fraction of unassigned transcripts as technical noise. In this study, we systematically investigated these unassigned RNAs (uRNAs), RNA molecules in in situ spatial transcriptomics data that are not assigned to segmented cells. We find that uRNA is widespread across spatial transcriptomics technologies and displays tissue-specific yet cross-technology consistent patterns. By quantifying the technical origin of these signals we identify that only a minor fraction arise from noise or diffusion. Importantly, while RNA diffusion has previously been proposed as a major contributor to mislocalization artifacts in iST data^28^, our analysis suggests that the spatial distribution of uRNA does not conform to classical diffusion patterns for most genes.

Our analysis also highlighted missegmentation as a major source of uRNA, even in datasets where membrane stains were used. Current state-of-the-art algorithms only partly alleviate this issue, which remains difficult to evaluate due to the lack of independent benchmarking studies. Missegmentation was often cell type-specific, reflecting the morphological complexity of certain cell types. Moreover, genes expressed by the same cells showed varying abundance in missegmented regions, with some transcripts even located far from their cell bodies. These findings challenge assumptions of intracellular homogeneity and spatial proximity in current segmentation algorithms, underscoring the need to incorporate such variability to improve performance.

For a substantial subset of genes, uRNA could not be explained by technical artifacts. The proportion of such non-technical uRNA varied across tissues and did not correlate with the expression of nearby segmented cells. In the brain, for instance, certain uRNA patterns are aligned with spatial and molecular expectations from RNA in cellular protrusions (e. g. axons, dendrites, glial processes). These structures often contain little RNA and, thus, are difficult to segment. To systematically analyze them, we proposed a strategy that infers protrusion-associated transcripts by assuming that unassigned reads originate from nearby cells expressing the same gene. While proven to be useful, this approach can underestimate or misattribute transcripts when cells downregulate genes in the soma or when protrusions extend far from the cell body, as in neurons or oligodendrocytes. Integration with spatial proteomics could improve accuracy by providing independent markers of cell identity, a possibility already supported by multimodal assays such as CosMx or CODEX.

A second potential biological source of uRNA is extracellular RNA (exRNA), which is actively secreted by cells in vesicles such as exosomes. While exRNA has been detected using the same chemistries employed in iST platforms^44^, its in situ detection remains a challenge. Vesicles typically carry low RNA content and are expected to diffuse through the tissue, making it difficult to distinguish them from other forms of uRNA. In our study, we were not able to resolve this contribution, but future efforts combining spatial transcriptomics with vesicle-specific staining could allow confident identification of vesicle-derived RNA.

In conclusion, our work provides the first systematic exploration of the multifaceted nature of unassigned RNA (uRNA) inspatial transcriptomics. By developing and applying troutpy, an open-source package for quantitative uRNA analysis, we provide a foundation for systematic exploration of these signals across tissues, technologies, and conditions. Beyond improving segmentation accuracy and quality control, uRNA analysis opens a window into subcellular transcript organization, cell-cell connectivity, and extracellular RNA biology. We anticipate that incorporating uRNA into future spatial workflows will refine how tissue architecture and cellular morphology are modeled in situ, turning what was once treated as noise into a new layer of biological insight.

## Code availability

Troutpy is a pip installable Python package, available at the following Github repository https://github.com/theislab/troutpy. Documentation is presented as a ReadTheDocs page and is available at https://troutpy.readthedocs.io/en/latest/. All code used to reproduce the analysis presented in this manuscript is available at the following Github repository: https://github.com/theislab/troutpy_reproducibility.

## Data availability

The data presented in this manuscript consists of a collection including datasets generated in previous studies^23-26^ and datasets released by the companies (i.e. 10X Genomics, Vizgen and Bruker) through their data release portals. Information regarding the characteristics of each dataset used through the manuscript, as well as links to their original publication and data source can be found in Extended Data Table 1. Furthermore, for reproducibility purposes, regions of interest of each dataset used through the study are available in Zenodo (10.5281/zenodo.17446348)

## Acknowledgements

This work was supported by the European Union (ERC, DeepCell - 101054957) and the DFG Leibniz Prize awarded to F.J.T. Additional support was provided by the Chan Zuckerberg Initiative Foundation (CZIF; grant CZIF2022-007488, Human Cell Atlas Data Ecosystem) and the Helmholtz Association through the joint research school “Munich School for Data Science - MUDS” (Grant number HIDSS-0006). F.D. was funded by the Hertie Network of Excellence in Clinical Neuroscience (5310272). S.M.S. was supported by the Vetenskapsrådet International Postdoc grant (2024-06623).

We would like to express our gratitude to Dr. Marco Grillo for his support and active discussions that contributed to improve the content of this work. We would also like to thank the members of the Theis lab for their discussions, feedback and support through the project. We would especially like to thank Lea Zimmermann for her helping in data retrieval and formatting and Philipp Angerer for his support in package building.

## Competing interests

F.J.T. consults for Immunai, CytoReason, BioTuring, Genbio and Valinor Industries, and has ownership interest in RN.AI Therapeutics, Dermagnostix, and Cellarity. R.K.R. advised iuvando GmbH and consults for RN.AI Therapeutics. S.M.S is cofounder of Spatialist AB. The other authors declare no conflict of interest.

## Online Methods

### Data collection

A total of 13 different datasets were compiled and used throughout this study from different sources, including previous publications^23-26^ and technology commercial partners (10X Genomics, Bruker and Vizgen). A summary of the datasets employed can be found in Extended Data Table 1. The collection includes datasets from different technologies and tissues, with panel sizes spanning from ~300 to 5000 genes. Details on how to access the datasets are provided in the Data Availability section.

### Data loaders & converters

Datasets from various in situ transcriptomics (iST) technologies were imported into the SpatialData (version 0.2.5) framework using the appropriate ‘spatialdata-io’ (version 0.1.5) loaders. Due to differences in file structure across technologies, custom data converters were developed to standardize each resulting SpatialData object into a common format. These converters are included within troutpy’s preprocessing submodule (e.g. troutpy.pp.xenium_converter). This harmonized organization ensures consistent data structure and reproducibility of the analysis.

### Cell segmentation and evaluation of segmentation impact

The analysis presented in the study relative to systematic characterization of uRNA in the corpus of 13 datasets spanning across tissues and technologies uses default multimodal segmentation provided by each platform as the basis for the analysis, as detailed in Figure 1A. For Xenium datasets where multimodal segmentation was not available, nuclei segmentation was used as default segmentation.

We also evaluated the impact of different segmentation strategies in the uRNA quantification. For this, we applied Proseg^14^ to both the mouse brain Xenium (5k) and the mouse brain CosMx regions of interest. The output of these segmentation approaches was processed using troutpy’s main workflow, including segmentation-free analysis using sainsc^27^ (version 0.2.1) and uRNA characterization using troutpy’s customized functions. The output of this analysis was compared to the default multimodal segmentation, as presented in Extended Data Figure 1.

### Cell expression processing

After cell segmentation, a cell-by-gene matrix with the transcriptomic composition of each identified cell was obtained. In order to identify the main cell populations present in the datasets analyzed, a common Scanpy-based processing workflow was followed. In summary, low quality cells, defined as cells where less than 30 counts of 10 different genes were detected, were filtered out. Library size-based normalization and log-transformation was then applied, prior to principal component analysis. After the construction of a k-nearest neighbors graph based on the selected principal components for each dataset (Code availability) cells were grouped using leiden clustering and represented in a low-dimensional space by applying Uniform Manifold Approximation and Projection (UMAP). Clustering and UMAP parameters, including resolution, were adjusted according to the expected populations in each dataset (Code Availability). Manual cell type annotations were performed as needed, based on identified differentially expressed genes, and incorporated into the analysis.

### Identifying cell-like signatures using segmentation-free analysis

To identify structures outside segmented cells that resemble the transcriptomic signatures of nearby cells, we applied segmentation-free analysis tools designed for spatial transcriptomics. Specifically, we adapted Sainsc^27^, a method that integrates a cell-segmentation-free approach with efficient processing of transcriptome-wide, nanometer-resolution spatial data. Sainsc generates high-resolution cell type maps with accurate cell type assignments and corresponding confidence scores, facilitating data interpretation. It is computationally efficient, scalable for interactive exploration, and compatible with common data analysis frameworks.

In our analysis, we first converted transcript localization data into a pixel-based representation, effectively binning transcripts into subcellular spatial units. Since troutpy processes individual transcripts, we needed to retrieve the transcript-to-pixel assignments, which is not directly available in the original Rust-based implementation of Sainsc. To enable this, we reimplemented part of Sainsc’s code in Python, allowing us to extract transcript-to-bin assignments for further analysis (Code availability).

To identify unsegmented regions that are transcriptomically similar to segmented cells, we first defined the mean expression signatures of cell populations identified based on segmented cells (see ‘Cell expression processing’ section). Using Sainsc, we computed the cosine similarity between intracellular expression signatures and each pixel. Pixels with high similarity to at least one intracellular signature were classified as cell-like structures.

To determine a similarity threshold, we compared the cosine similarity of pixels from segmented cells with that of pixels from unassigned regions. We reasoned that unassigned pixels with similarity values within the range observed for segmented cells likely represent “cell-like” structures, whereas pixels with lower similarity reflect non cell-like regions (labeled as “other”). We defined the threshold as the 10th percentile of the cosine similarity values of segmented-cell bins. This choice ensures that ≥90% of bins from true cells exceed the threshold, while allowing for up to 10% possible missegmentation. Unassigned pixels with a maximum similarity above this threshold were classified as cell-like, and the rest as other.

#### Evaluating the impact of Sainsc hyperparameters on uRNA quantification

The analysis presented in our manuscript is based on the identification of cell-like regions. As exposed in the previous section, this is done by adapting Sainsc to distinguish regions that transcriptomically resemble segmented cells from the ones that do not. Since both the Sainsc algorithm and the workflow where it is implemented are dependent on given hyperparameters, we explored the impact of tuning these hyperparameters in the characterization of unassigned RNA. For this, we applied multiple times the Sainsc workflow to the Xenium mouse brain region of interest, used through the manuscript, trying all possible combinations of the hyperparameters. Hyperparameters tested include percentile threshold (10, 20), bin size (2,5,10), background filter (0.2,0.4) and gaussian kernel (0,2,4). For each run, uRNA was classified between “cell-like” and “other”. To compare the runs, the transcriptomic composition of “other” uRNA pools was compared using cosine similarity. In addition, we also compared the proportions of uRNA identified as “cell-like” across runs. Our analysis revealed a high consistency in the uRNA composition across all of the parameters used.

#### Exploration of uRNA corresponding to cell-like regions

To investigate the nature of cell-like uRNA regions identified by Sainsc, we analyzed data from the Xenium Prime mouse brain dataset. Sainsc assigns each pixel to the cell type with the highest cosine similarity to predefined cell type expression signatures. Using this assignment, we focused only on cell-like regions (as defined in previous sections) and, for each cell type, calculated the fraction of transcripts located in these regions relative to the total RNA (transcripts inside segmented cells plus those in cell-like regions).

To assess the spatial distribution of cell-like regions, we measured the minimum distance between each transcript in these regions and the nearest transcript from a segmented cell of the same Sainsc-assigned cell type. We classified distant cell-like uRNA as transcripts located more than 10 µm away from any segmented-cell transcript of the same type. Results are shown in Extended Data Figure 2.

### Aggregation of unassigned RNA

Unassigned transcripts are typically defined in iST datasets based on their position (either 2D or 3D) and their gene identity. With the aim of either analyzing the local context of each unassigned transcript or to speed up some tasks, it can be useful to aggregate unassigned transcripts into bins, or pixels, that summarize the local uRNA composition of a certain tissue area. This would be analogous to cell segmentation, where we group transcripts based on cellular boundaries, generating an expression vector. In our work, this task is performed using the function ‘troutpy.pp.aggregate_urna’. While this function supports multiple strategies to aggregate unassigned transcripts (see troutpy’s documentation), the analyses presented through this study are based on the ‘bin’ strategy.

In this approach, the tissue plane is tessellated into a regular grid of fixed-size squares that cover the full spatial extent of the transcript coordinates. Each square is represented as a polygon geometry, and its centroid is stored to serve as the spatial coordinate of the aggregated unit. Transcripts located outside segmented cells are then assigned to their corresponding grid square, and gene-level counts are aggregated within each bin. The result is a segmentation-free expression table, analogous to a cell-level AnnData object, where each row corresponds to one bin and each column to a gene. Square sizes, representing the length of each square, were set to 10 micrometers through all the analyses presented in the manuscript, unless specified otherwise.

### Unassigned RNA characterization

Several metrics were developed to characterize the uRNA nature of each gene profiled. In the following sections, we detail the different metrics used in the analysis of the datasets presented through the manuscript.

#### Quantification of transcripts overdetected outside segmented cells

To determine whether unassigned RNA expression exceeds background noise, we developed a statistical framework that compares gene-associated transcript counts to non-gene control features. These control features (predefined codewords) represent background signals. We first calculated the distribution of counts for these controls and defined the 99th percentile as the noise threshold.

When no control probe is available, we instead inferred noise by analyzing the spatial distribution of transcripts inside versus outside segmented cells. Specifically, we computed the ratio of transcript counts overlapping segmented cell regions relative to counts outside cells, normalized by their respective areas. Genes for which the normalized count outside cells exceeds that inside cells are flagged as potentially noisy. We then use the median total count of these flagged genes to define a noise threshold. For each gene, we then assessed overexpression by measuring the fold enrichment of its unassigned counts relative to this threshold, allowing us to identify transcripts with significantly elevated expression outside segmented cells. Log fold change values were calculated to quantify the extent of deviation from background expression:

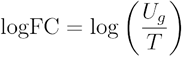

where *U*_*g*_ is the observed uRNA count for gene *g* and *T* is the percentile-based noise threshold, either derived from control features or computed. Overall, these metrics enable the downstream identification of genes with enriched extracellular expression, providing insights into transcript distribution and potential biological relevance.

#### Identification of spatially variable uRNAs

To assess whether unassigned transcripts for each gene exhibited spatial variability, we computed spatial autocorrelation on pre-binned unassigned transcript counts (see *Aggregation of unassigned RNA* section). Our strategy was built on top of Squidpy’s (version 1.6.2) implementation of Moran’s I, which measures the degree of spatial clustering in a gene’s expression pattern. This approach consists of two steps: (i) spatial neighbors of the precomputed bins were identified using ‘squidpy.gr.spatial_neighbors’, and (ii) Moran’s I was calculated with ‘squidpy.gr.spatial_autocorr’. Each gene received a Moran’s I score between −1 and 1. Values approaching 1 indicated strong positive spatial autocorrelation, consistent with spatially variable expression, while values near −1 denoted negative spatial autocorrelation, where high and low expression alternate in a checkerboard-like pattern.

#### Quantification of proportion of unassigned RNA for each gene

To quantify unassigned RNA for each gene, we computed the proportion of transcripts located outside segmented cells using the ‘troutpy.tl.extracellular_enrichment’ function within troutpy. In this context, the term “extracellular” is a simplified term used to refer to all unassigned RNA, in contrast with intracellular RNA, referring to RNA segmented into cells. Essentially, the function calculates per-gene proportions of unassigned and intracellular transcripts, along with the log fold change between them.

#### Quantification of intracellular-uRNA local correlation

The correlation between the intracellular RNA expression and its surrounding uRNA, for each gene, is calculated using ‘troutpy.tl.in_out_correlation’. Essentially, this function computes, for each gene, the Spearman correlation between the gene expression measured across cells and each gene’s abundance found in spatially proximal unassigned RNA bins. Therefore, unassigned RNA bins are generated prior to this step by aggregating transcripts located outside segmented cells into fixed-area spatial units (see *‘Aggregation of unassigned RNA’* section).

For each cell *i* and gene *g*, the expression from the *k* nearest unassigned RNA bins was averaged as:

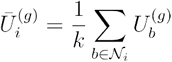

where *𝒩 i* denotes the nearest unassigned RNA bins to cell *i*, and 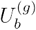 is the expression of gene in bin *b*. In the analysis presented through the manuscript, the number of neighbors *k* was adjusted per dataset to approximate the mean number of transcripts per cell. Specifically, *k* was estimated by comparing the typical transcript abundance in segmented cells and unassigned RNA bins, using the ratio of their median counts. Finally, Spearman’s correlation was then computed between the aggregated unassigned RNA expression 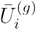 and the intracellular expression 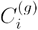 across all cells.

### Multivariate analysis of uRNA organization

After characterizing gene-specific uRNA patterns across datasets and exploring their potential origin, we focused on characterizing the multivariate nature of uRNA. In this manuscript, multivariate analysis is only presented for Xenium’s multimodal mouse brain (Figure 4, Extended Data Figure 5) and human femur datasets (Extended Data Figure 6). For this, we first grouped uRNA into spatial bins based on the spatial position of individual transcripts (see ‘Aggregation of unassigned RNA’ section). Bins were then filtered based on a minimum number of counts, transformed and clustered based on gene expression profiles, using the same strategy followed for segmented cells (see ‘Cell expression processing’ section). In the mouse brain dataset, clusters obtained were annotated based on gene expression profile and spatial localization. Since uRNA clusters mapped accurately diverse brain structures, cluster annotation was done based on the brain structures defined in the Allen Mouse Brain Atlas^45^.

#### DRVI analysis

With the aim of disentangling the molecular complexity of predefined uRNA bins, we applied DRVI^38^ (version 0.1.9), which enables learning interpretable disentangled dimensions. For each of the 20 non-vanished DRs identified in the mouse brain dataset, we generated both spatial maps and gene contribution bar plots. Furthermore, since this variability is hypothesized to emerge from segmented cells, we also use the function ‘troutpy.tl.factors_to_cells’ to compute a DR score for each profiled cell. For this, we defined for each DR a contribution vector for summarizing the contribution of each gene to dimension. Each cell’s projected score for each DR was then achieved by the multiplication of the contribution vector by the expression of each cell. As a result, a DR-specific projected score was achieved for each DR across all segmented cells.

#### Non-negative matrix factorization (NMF)

Besides DRVI, other disentanglement approaches are included in troutpy including Latent Discriminant Analysis (LDA)^46^ and non-negative matrix factorization (NMF)^47^. NMF is indeed used in the manuscript to identify gene expression programs contributing to the uRNA pool in both the mouse brain Xenium dataset (Extended Data Figure 5) and in the Xenium femur dataset (Extended Data Figure 6). After grouping uRNA into spatial bins based on the spatial position of individual transcripts (see ‘Aggregation of unassigned RNA’ section), NMF was applied to identify 10 factors driving the uRNA variability. For each factor, spatial maps based on the factor score of each spatial bin, as well gene loadings were visualized. Based on the gene loadings’ vector obtained for each factor, factor scores were calculated for segmented cells based on their expression profile, as done in DRVI analysis. Gene set enrichment analysis (GSEA) was performed using gseapy (1.1.9) by selecting all genes contributing more than 1% to each factor’s total loading. The enrichment of these genes was assessed against the *GO_Cellular_Component_2021* gene set database for mice, using the Enrichr implementation of GSEA.

#### Staining quantification

Since spatial platforms provide morphological staining for segmentation, we aimed to compare whether the high factors/dimensions identified using DRVI or NMF could be linked to a difference in the intensity signal of these stainings. To achieve this, we computed, for each precomputed uRNA bin, the mean intensity in the k-nearest image pixels (k = 10), across morphological stainings. After obtaining mean intensity values for each morphological staining across uRNA bins, we transformed the raw intensities into z-scores. This was done by centering the values for each staining around the average intensity across all uRNA bins and scaling them by the corresponding standard deviation. These z-scores indicate the deviation in signal intensity from the distribution of intensities obtained, for a certain staining, across uRNA bins. These z-scores indicate the deviation in signal intensity from the intensities obtained, for a certain staining, across uRNA bins. Therefore, positive values represent a higher intensity than expected by random, whereas negative values represent lower intensities than expected by random. We visualized the relation between factors’ scores and staining intensities by visualizing, for each factor, the mean z-score found in the 10% most positive uRNA bins. In DRVI analysis, depending on the direction of the dimension, either the 10% most positive or negative cells for each dimension were used.

### Computation of cell centric-based uRNA scores

#### Identification of uRNA’s source cells and uRNA source score

To identify the most probable source of each unassigned RNA (uRNA) transcript, we computed source scores that combine spatial proximity with gene expression levels across candidate cells. This approach computes the probability of a cell to be the source of a specific uRNA transcript based on its distance to the transcript and the expression of its corresponding gene.

For each unassigned transcript *t*_*j*_ associated with gene *k*, we first identified all cells with non-zero expression of gene *k*. The Euclidean distance between the position of transcript *t*_*j*_ and the centroid of each candidate cell *c*_*i*_ was computed as:

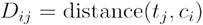

A weighted score *S*_*ij*_ was then assigned to each transcript-cell pair using the following exponential decay function:

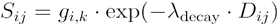

where *g*_*i,k*_ is the expression level of gene *k* in cell *c*_*i*_, and λ_dencay_ is a user-defined decay constant (in our analysis, λ_dencay_ = 0.1) that controls how rapidly the score decreases with increasing distance. This formulation gives higher scores to nearby cells that express the relevant gene.

To estimate the contribution of each cell type, the scores were aggregated across all cells of the same type. The total source score for transcript *t*_*j*_ with respect to cell type *ct*_*s*_ was computed as:

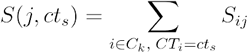

where *C*_*k*_ is the set of all cells expressing gene *k*, and *CT*_*i*_ denotes the cell type of cell *c*_*i*_. The result is a raw score indicating the total proximity-weighted expression for each cell type. To interpret these scores as relative probabilities, we normalized them across all cell types for each transcript. A small residual constant *ε* =10 ^− 6^was added for numerical stability, yielding:

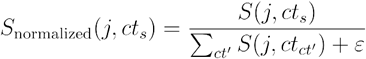

This produces a distribution over cell types for each transcript, representing the relative likelihood that it originated from each type.

In addition to transcript-level inference, we also computed a cell-level source score by summing the contributions from all extracellular transcripts, considering both proximity and gene expression. The source score for cell cjc_j was defined as:

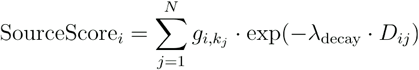

where *k*_*j*_ is the gene associated with *t*_*j*_, and *D*_*ij*_ is the distance from transcript *t*_*j*_ to cell *c*_*i*_. Only transcripts for which cell expresses the relevant gene were included.

#### Identification of uRNA’s target cells

To infer the most likely target cell types for each unassigned RNA (uRNA), we computed proximity-based scores. These target scores are calculated using spatial distance and aggregated by annotated cell types to capture cell type-specific proximity patterns.

For each extracellular transcript *t*_*j*_, we calculated the Euclidean distance *D*_*ij*_ between the transcript’s spatial coordinates and the centroids of the *k* nearest cells *c*_*i*_ identified using a k-nearest neighbors (kNN) search in two-dimensional space. Each transcript-cell pair was then assigned a proximity score *T*_*ij*_ based on an exponential decay function:

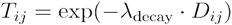

Here, λ_dencay_ is a tunable decay parameter that controls how strongly the score decreases with distance. As it occurred with source score calculation, λ_dencay_ was also set to 0.1 through the analysis presented in the manuscript. This formulation gives higher scores to cells that are spatially closer to the transcript, reflecting a higher likelihood of interaction or molecular targeting.

To capture cell type-specific interactions, we aggregated the proximity scores for all nearby cells belonging to the same cell type *Ct*_*t*_. The total score for transcript *t*_*j*_ and cell type *Ct*_*t*_ is given by:

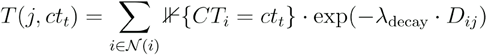

where 𝒩 (*i*) denotes the set of the *k* nearest neighboring cells to transcript *t*_*j*_, and *CT*_*i*_ is the annotated cell type of cell *c*_*i*_.

To facilitate interpretation and comparison across transcripts, the target scores for each transcript were normalized across all cell types to sum to 1:

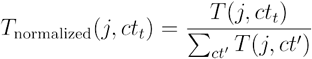

This normalization yields a probability-like distribution for each transcript based on spatial proximity.

#### Quantification of uRNA-mediated cell-cell contacts

To assess whether unassigned RNA (uRNA) captures cellular protrusions and, in turn, cell-cell contacts that remain undetected by standard segmentation, we quantified interactions at two levels: (i) direct spatial contacts between segmented cells, and (ii) extended contacts revealed through uRNA source-target inference. We first considered only segmented cells in the dataset. Each cell *c*_*i*_ is represented by its centroid coordinates *X*_*i*_ ∈ ℝ^*d*^ and associated with a cell type label *t*_*i*_ ∈ *T*, where *T* is the set of annotated cell types.

For a given distance threshold *r*, the spatial neighborhood of a cell *c*_*i*_ is defined as:

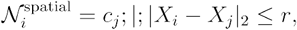

where ∣ · ∣ _2_ denotes the Euclidean distance. Spatial neighborhoods were computed efficiently using a BallTree algorithm.

From these neighborhoods we constructed a cell-cell contact matrix 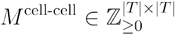, where each entry counts the number of contacts between pairs of cell types:

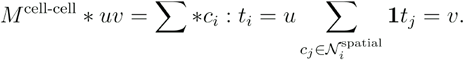

This matrix provides a baseline representation of spatial contacts detectable from cell centroids alone.

Next, we incorporated uRNA source-target assignments (see previous sections) to extend the neighborhood definition. Let *S*=(*s*_*k*_, *t*_*k*_, *d*_*k*_) denote the set of inferred source-target transcript pairs (see sections above), where *s*_*k*_ is the source cell, *t*_*k*_ is the target cell, and *d*_*k*_ is the distance between them. We filtered these pairs to retain only additional links not already captured by the direct spatial neighborhood:

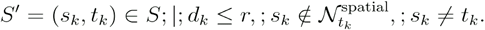

The extended neighborhood of cell was then defined as:

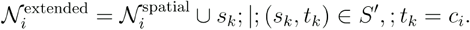

Importantly, only cells presenting a number of source-target pairs above threshold *k*_*t*_ where considered. For the analysis presented in the study, a minimum of 10 uRNA molecules suggesting a source-target relation between pairs of cells *k*_*t*_ >10 were considered. Using the extended neighborhood, we computed a combined contact matrix:

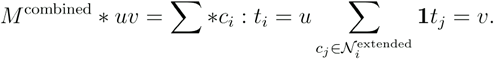

Finally, to isolate the contribution of extracellular RNA, we derived a uRNA-specific contact matrix:

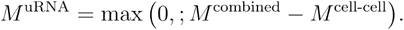

This captures interactions uniquely attributable to uRNA-mediated proximity, excluding those already present in the direct spatial contact matrix.

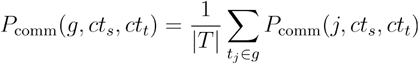

#### Calculation of uRNA contribution score

To quantify how much each cell contributes to the unassigned RNA (uRNA) pool, we defined a uRNA contribution score that combines gene-level uRNA abundance with the distribution of gene expression across cells. This score reflects the hypothesis that a gene’s unassigned signal originates proportionally from the cells that express it.

For each gene *g*, the total number of uRNA transcripts detected outside cells is denoted *U*_*g*_. For each cell, we use the raw gene expression matrix *X*, where *X* _*c,g*_ represents the count of gene *g* in cell *c*. A gene is considered expressed in a cell if *X* _*c,g*_>0.

We define the uRNA contribution score for cell as:

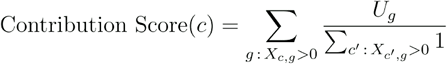

That is, for every gene expressed by a cell, we distribute the extracellular count *U*_*g*_.evenly across all cells expressing that gene. The score for a given cell is the sum of its weighted contributions across all genes it expresses. To control for differences in total RNA abundance across cells, we also compute a normalized uRNA contribution score, defined as:

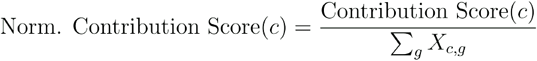

This normalized score captures the extent to which a cell contributes to the uRNA pool relative to its total intracellular transcript count.

#### Assessing RNA diffusion

To evaluate whether unassigned RNA exhibit patterns consistent with passive diffusion, we modeled their displacement distributions relative to their precomputed source cells (see above). For each gene, the displacement distances *r* of unassigned transcripts were collected. Genes with fewer than 10 unassigned transcripts or with no measurable displacement were excluded from analysis.

The observed distance distribution was compared against the Rayleigh distribution, which describes the probability density of radial displacements under two-dimensional isotropic diffusion:

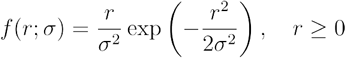

where is σ the scale parameter estimated by maximum likelihood.

To assess whether transcript displacements follow Rayleigh statistics, we applied Kolmogorov-Smirnov test (KS) comparing the measured distribution and the Rayleigh distribution fitted to each gene’s distances, assessing significance by applying a bonferroni correction based on the number of genes tested in each dataset. Global KS-test comparing the measured distribution and the Rayleigh fitted distribution for all uRNA molecules per sample was also computed.

Under a simple two-dimensional diffusion process, the mean squared displacement (MSD) is related to the diffusion coefficient *D*:

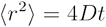

For the Rayleigh distribution, the MSD is:

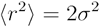

Thus, the effective diffusion coefficient can be estimated as:

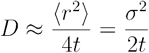

where *t* represents the characteristic timescale between RNA release and measurement (unknown but constant across genes). In practice, we report the mean displacement ⟨*r* ⟩.as a relative proxy for diffusion strength across genes:

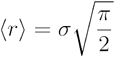

#### Simulation of RNA diffusion

To model evaluate the capacity of our strategy developed to capture diffusion patterns in iST datasets, we developed a simulation framework that generates synthetic datasets of cells and corresponding diffused RNA.

A population of a predefined number of cells was randomly positioned within a 2D spatial domain of size (1000 × 1000) units, sampled from a uniform distribution. Each cell was assigned an expression profile for a defined number of genes, with transcript counts drawn from a Poisson distribution (λ=20) to mimic variability in intracellular expression. For each gene in each cell, a fraction of transcripts was designated as unassigned according to a release parameter (exRNA_factor). The resulting number of unassigned molecules (uRNA) was computed as the integer product of intracellular abundance and the release parameter. uRNAs were spatially displaced from their source cell according to a 2D Rayleigh diffusion model. Specifically, each molecule was assigned a radial displacement sampled from a Rayleigh distribution, defined by a scale parameter (scale_rayleigh), and an angle sampled uniformly from [0, 2π). This generates a circular diffusion profile centered at the cell, with the majority of molecules concentrated within approximately one scale_rayleigh unit of the source.

To evaluate if our strategy to capture diffusion patterns could correctly characterize these patterns, we then applied our approach to the simulated datasets. In short, for each gene, the empirical distribution of uRNA distances from the source cell was compared against the theoretical Rayleigh distribution using multiple statistical tests. Parameters of the Rayleigh distribution were fitted to the observed distances via maximum likelihood. Goodness-of-fit was assessed with the Kolmogorov-Smirnov (KS) test, comparing Rayleigh fits to kernel density estimates. Genes with KS test p-values below a significance threshold (ks_thresh=0.05) were classified as “non-diffusive.” The main simulation output is the proportion of genes deviating from Rayleigh diffusion under each parameter setting. Simulations were systematically performed across combinations of hyperparameters, varying the number of cells (n_cells ∈ {50, 100, 200, 400}), diffusion scales (scale_rayleigh ∈ {5, 10, 20, 50}), and release fractions (exRNA_factor ∈ {0.05, 0.1, 0.2}).

### Extrasomatic RNA quantification

#### Evaluating the extrasomatic RNA quantification in nuclear/cytoplasmic data

To evaluate the accuracy of troutpy in quantifying extrasomatic RNA, we used the Xenium prime mouse brain dataset used through the manuscript. Since the evaluation of the strategy requires a ground truth and we can’t have this information for the uRNA, we focused on subcellular compartmentalization. Specifically, transcripts localized to the cytoplasm were treated as extrasomatic, whereas nuclear transcripts were considered intracellular. To ensure a direct comparison, we excluded all transcripts not assigned to cells (uRNA) and redefined cell-associated expression as consisting only of nuclear-localized transcripts, while cytoplasmic transcripts were treated as unassigned. We then applied ‘troutpy.tl.urna_source_score’ to estimate the likely cell of origin for each cytoplasmic transcript. This function uses an exponential distance-weighted model informed by nearby nuclear expression. Transcript-level source probabilities were aggregated by gene and cell type to obtain predicted extrasomatic proportions. Finally, we compared the predicted values to the true extranuclear proportions, calculated as cytoplasmic / (cytoplasmic + nuclear) per gene and cell type. Agreement was quantified using both the Pearson correlation coefficient (PCC) and cosine similarity.

### Use of Large Language Models

Large Language Models (LLMs), specifically OpenAI’s ChatGPT (GPT-5.1, 2025), were used during the preparation of this manuscript to assist with rewording, improving coherence and readability, and correcting grammar and phrasing. All content was reviewed and validated by the authors to ensure accuracy and scientific integrity.

## Extended Data Figures

**Extended Data Figure 1.**
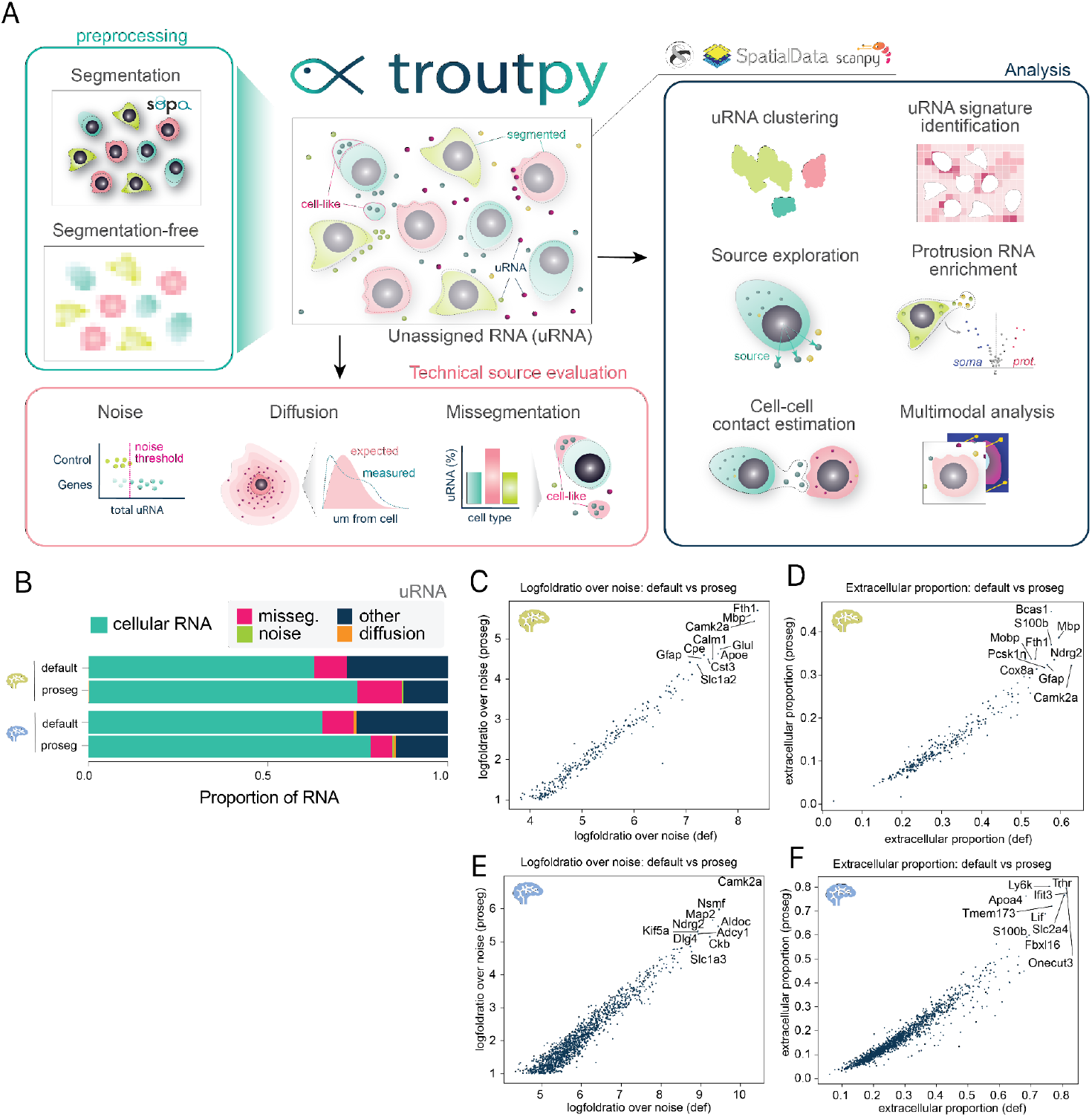
Description of troutpy and evaluation of alternative segmentation strategies. **(A)** Schematic representation of the main functionalities supported by troutpy, a python package developed to analyze unassigned RNA in iST datasets. **(B)** Barplot comparing the proportion of transcript assigned and unassigned to cells in both the CosMx and Xenium mouse brain datasets when using default segmentation and transcript-based Proseg segmentation. Unassigned RNA is colored depending on whether it can be attributed to missegmentation, noise, diffusion or none of the previous.(**C)** Scatter plot representing the uRNA abundance, represented as log fold ratio over noise threshold, identified the CosMx mouse brain dataset when using default segmentation (x-axis) in contrast with proseg segmentation (y-axis). **(D)** Scatter plot representing the uRNA proportion identified the CosMx mouse brain dataset when using default segmentation (x-axis) in contrast with proseg segmentation (y-axis). **(E)** Scatter plot representing the uRNA abundance, represented as log fold ratio over noise threshold, identified the Xenium mouse brain dataset when using default segmentation (x-axis) in contrast with proseg segmentation (y-axis). **(F)** Scatter plot representing the uRNA proportion identified the Xenium mouse brain dataset when using default segmentation (x-axis) in contrast with proseg segmentation (y-axis).

**Extended Data Figure 2.**
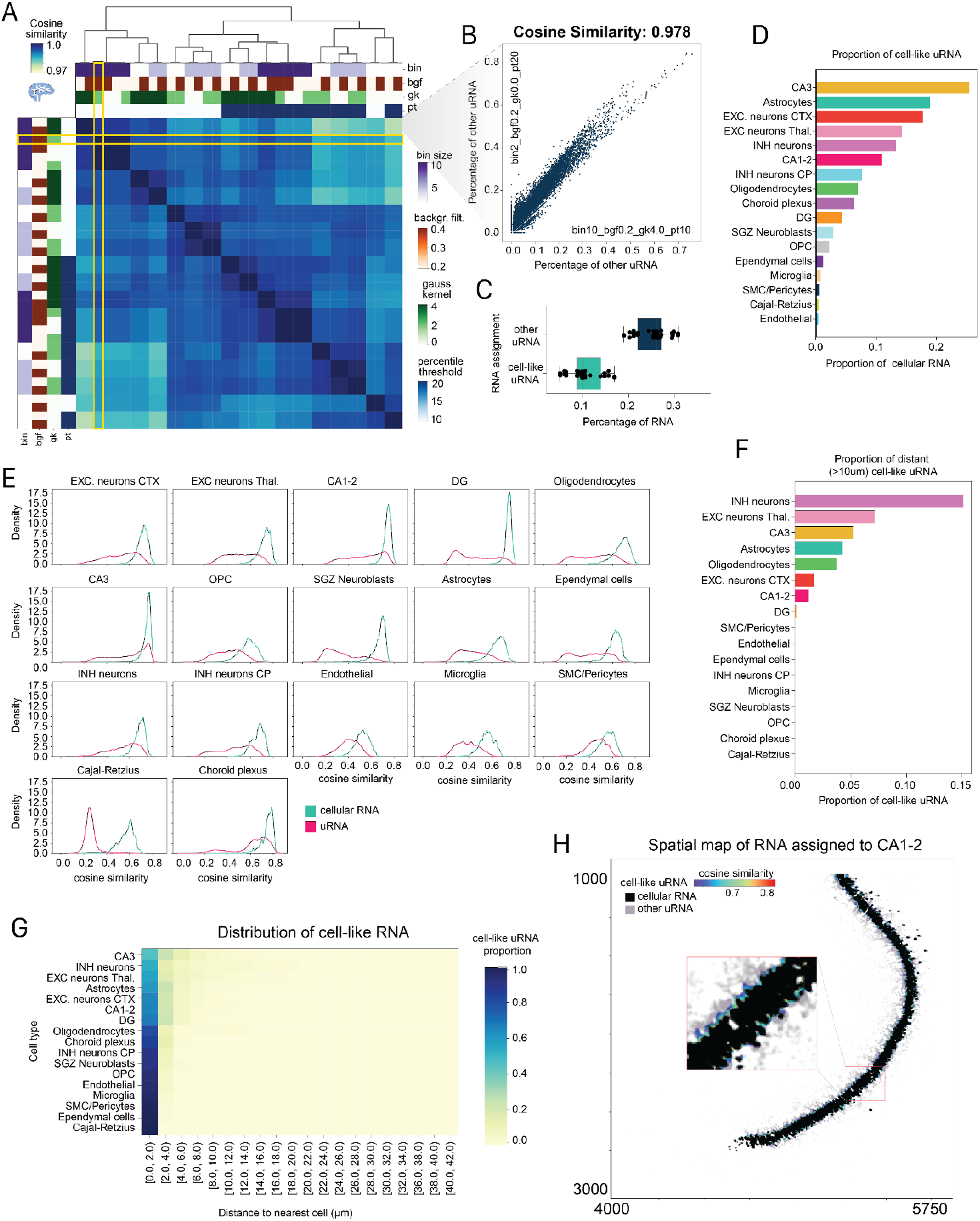
Characterization of cell-like uRNA region in Xenium mouse brain dataset. **(A)** Heatmap showing the cosine similarity between the percentage of uRNA classified as “other” (i.e., not defined as cell-like by Sainsc) for each gene across multiple runs of the method with different parameter settings. Tuned parameters included bin size, background filter, Gaussian kernel, and percentile threshold. The yellow ribbon marks the default settings used throughout the manuscript. **(B)** Scatter plot comparing, for each gene, the percentage of uRNA classified as “other” for each gene in the two Sainsc runs that showed the lowest cosine similarity (as defined in panel A). **(C)** Box plot representing, for all the Sainsc runs presented in A, the percentage of total RNA found to be cell-like uRNA and other uRNA. Individual points represent the proportions identified by individual runs. **(D)** Bar plot showing, for each cell type, the proportion of RNA corresponding to cell-like structures that were not originally captured by segmentation, as identified by Sainsc. **(E)** Density plots of cosine similarity between local transcriptomic regions defined by Sainsc and predefined cell type signatures. Each region was assigned to the cell type with the highest similarity score. For regions assigned to each cell type, we show the distribution of cosine similarity values, distinguishing between regions originally located within segmented cells (cellular RNA) and those outside cells (uRNA). **(F)** Bar plot showing, for each cell type, the proportion of cell-like uRNA found at least 10 micrometers away from any presegmented cell. **(G)** Heatmap showing the distribution of distances (in 2 µm bins, x-axis) from cell-like regions to the nearest segmented cell, stratified by the cell type assigned to each region (y-axis). Values are normalized to illustrate the relative proportion of cell-like regions for each cell type across distance ranges. **(H)** Spatial map of transcripts assigned to CA1-2 cells by Sainsc. Transcripts within segmented cells (cellular RNA) are shown in black. Among uRNA, those classified as cell-like (cosine similarity to the CA1-2 reference above the threshold) are colored by similarity score according to the colormap, while those below the threshold are shown in grey. Coordinates are in micrometers.

**Extended Data Figure 3.**
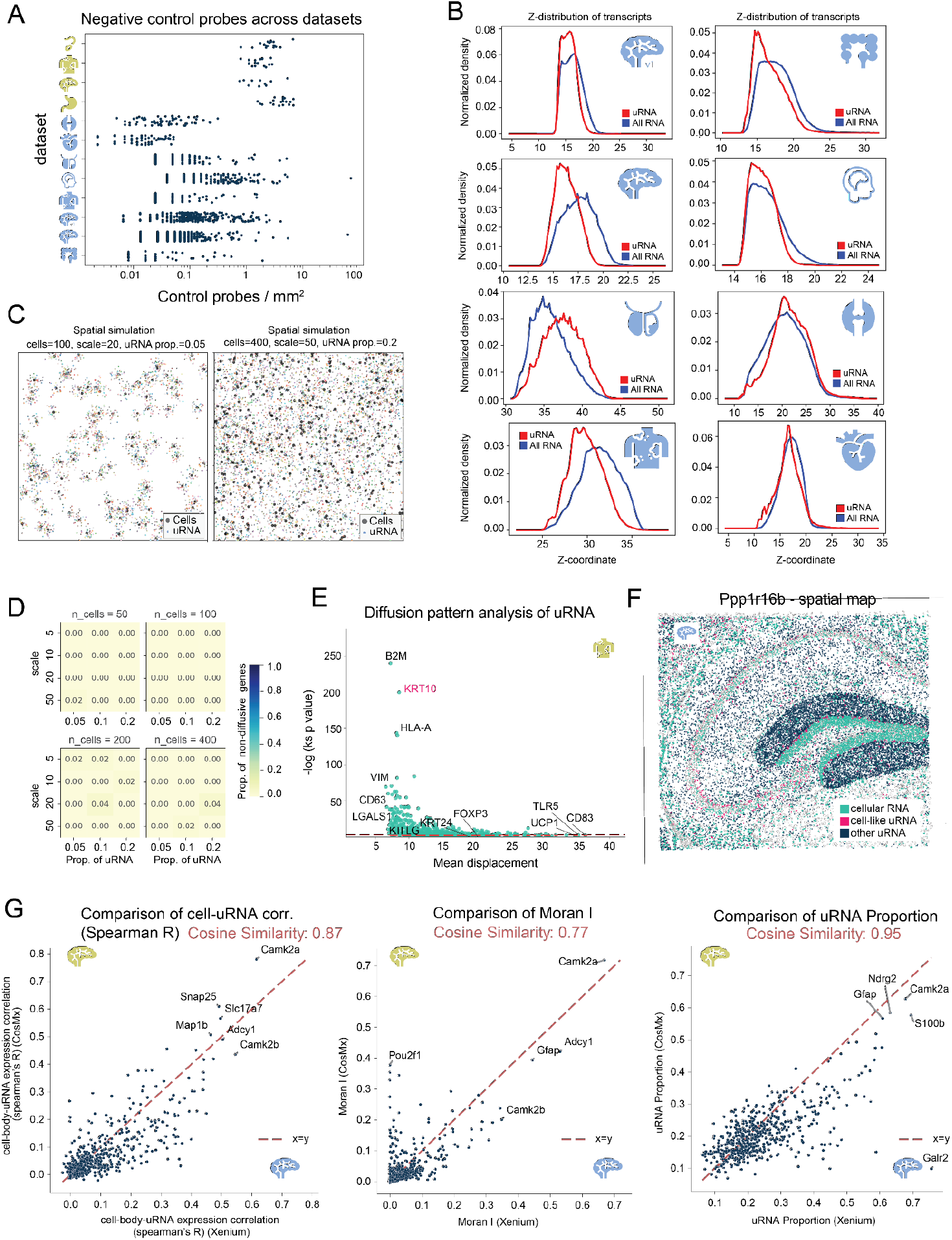
Exploration of the technical sources and consistency of uRNA. **(A)** Strip plot representing, for each dataset analyzed that included control probes (CosMx & Xenium), the amount of transcripts that each control probe presented per square millimeter. Individual dots present individual control probes. **(B)** Distribution of the transcripts across the Z-axis in the Xenium regions of interest analyzed, one subplot per dataset. Distribution of both uRNA and all transcripts profiled are represented. **(C)** Spatial maps of simulated datasets used to evaluate the strategy for identifying diffusion patterns. Spatial coordinates span a 1000 × 1000 space. Two examples of simulations generated from predefined parameters are shown. **(D)** Heatmaps showing the proportion of genes classified as “non-diffusive” across simulated datasets. Each heatmap corresponds to a different number of cells. The x-axis indicates the proportion of unassigned RNA, and the y-axis indicates the scale parameter of the Rayleigh distribution. **(E)** In the scatter plot (right), each point represents a gene, with the x-axis showing the mean displacement distance of its transcripts from their source (in micrometers), and the y-axis showing the −log_10_(p-value) from a Kolmogorov-Smirnov test comparing the observed distribution to a fitted Rayleigh distribution. The horizontal line indicates the significance threshold (p = 0.05); genes above this line are considered inconsistent with a diffusion-like distribution. **(F)** Spatial map of the Xenium mouse brain 5k sample. Transcripts assigned to Ppp1r16b are displayed and colored by RNA status: cellular RNA, or uRNA. uRNA is further subdivided based on spatial context into cell-like uRNA (found in cell-like regions) and other uRNA. Cell boundaries from the default Xenium multimodal segmentation are overlaid in the background. **(G)** Scatter plot comparing gene-level uRNA proportion (right), spatial variability (mid) and cell-uRNA correlation (left) between biologically comparable hippocampal regions in the Xenium (x-axis) and CosMx (y-axis) mouse brain samples (left). A diagonal reference line (x = y) is included to aid visual comparison. The names of the top five genes with the highest uRNA abundance in both samples are labeled on each plot.

**Extended Data Figure 4.**
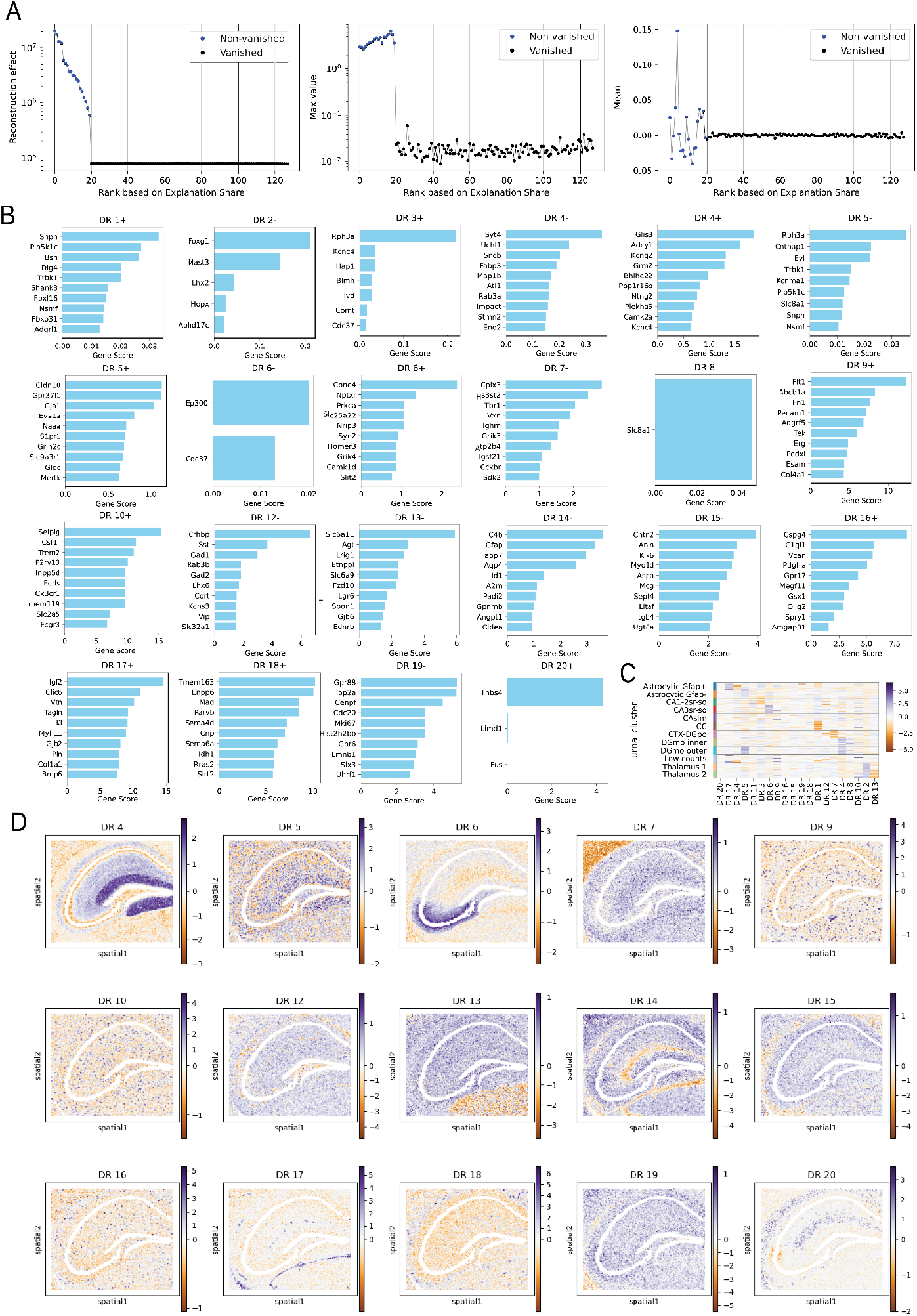
Expanded DRVI analysis using Xenium mouse brain’s unassigned RNA reveals cell type and region-associated gene programs. **(A)** Scatter plots indicating, for each DR (x-axis), its reconstruction effect, max and mean value (y-axis). DRs are colored based on whether they are found to be vanished or non-vanished. **(B)** Bar plot representing, for each DR, the contribution of the genes contributing the most to each DR both on their positive (+) and their negative (−) space. Genes with a contribution above 0 are represented, up to a maximum of 10 genes per plot. DRs whose contribution is presented in Figure 4D are not represented. **(C)** Heatmap representing the DR scores for each uRNA bin, grouped by their uRNA cluster. **(D)** Spatial map of the uRNA bins, colored by their score for each DR, according to the DRVI analysis performed. Only DRs not shown in Figure 4D are represented.

**Extended Data Figure 5.**
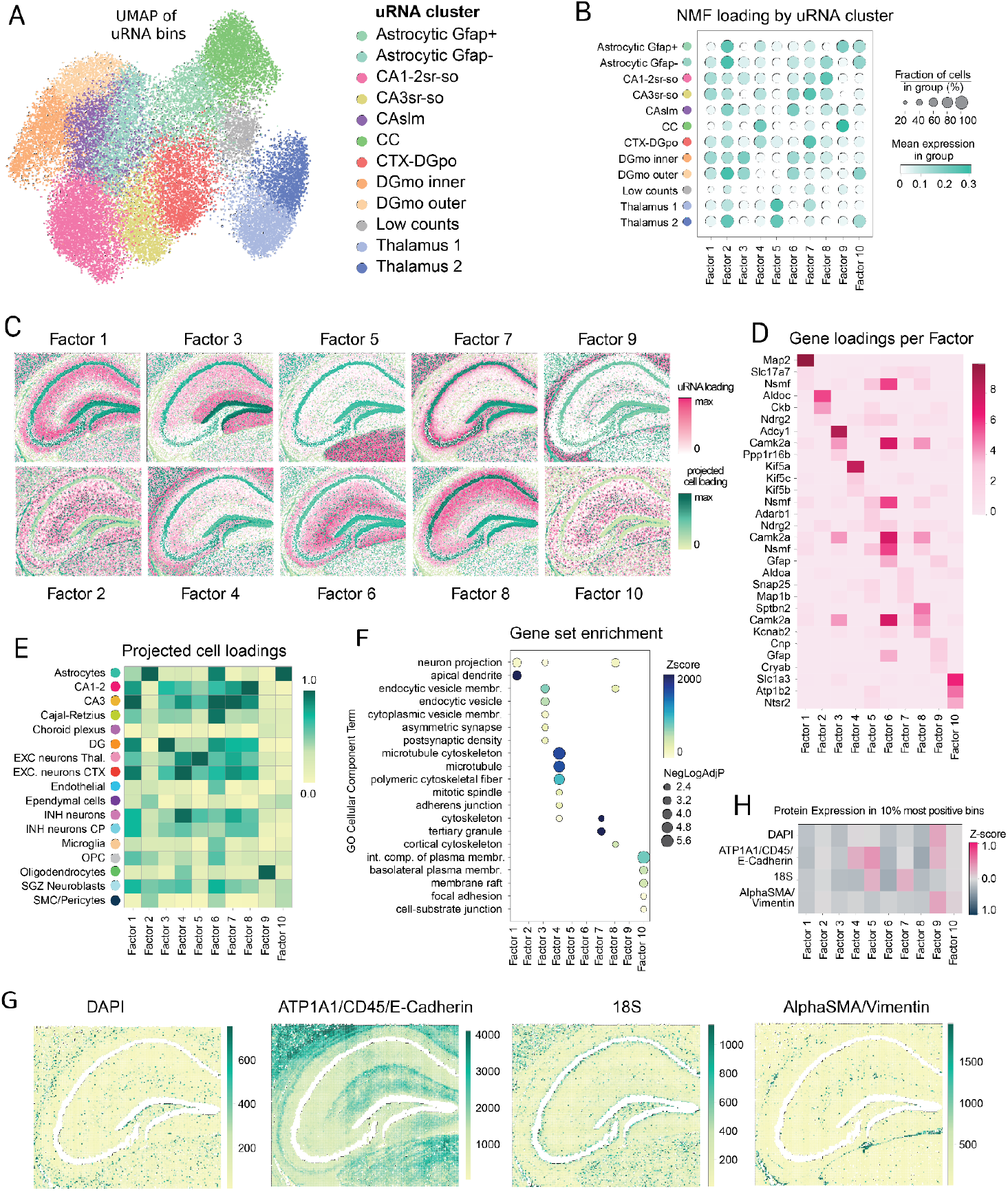
NMF analysis performed using Xenium mouse brain’s unassigned RNA highlights spatial and morphological differences in the RNA composition outside segmented cells. **(A)** UMAP of the uRNA bins identified in the Xenium mouse brain dataset, colored by uRNA cluster. **(B)** Dot plot illustrating the cell loading of each factor obtained from NMF analysis for the different uRNA clusters annotated in A. Dot color indicates the mean cell loading, whereas dot size represents the percentage of bins positive with loading different to zero for each given factor. **(C)** Spatial maps showing the factor scores for each uRNA bin across all NMF-derived factors (pink scale). Segmented cells are overlaid and colored according to their corresponding projected factor scores (green scale). **(D)** Heatmap showing the top three genes with the highest loadings for each factor, obtained from NMF-analysis shown in D, displaying their gene loading values across all factors. **(E)** Heatmap representing the mean projected cell loadings of each NMF factor on each of the cell types obtained from the annotation of segmented cells, normalized by factor. **(F)** Dot plot showing GO Cellular Component enrichment (GSEA) of the top-scoring genes for each factor. Only significant terms are displayed. Dot color indicates the z-score, and dot size reflects the −log_10_ of the adjusted p-value. **(G)** Heatmap showing the Z-score of staining intensities (y-axis) across complementary Xenium stainings, calculated within the top 10% most positive bins for each factor (x-axis). **(H)** Spatial maps of uRNA bins, colored by the intensity of the morphological stainings provided in the Xenium output at the location of each bin.

**Extended Data Figure 6.**
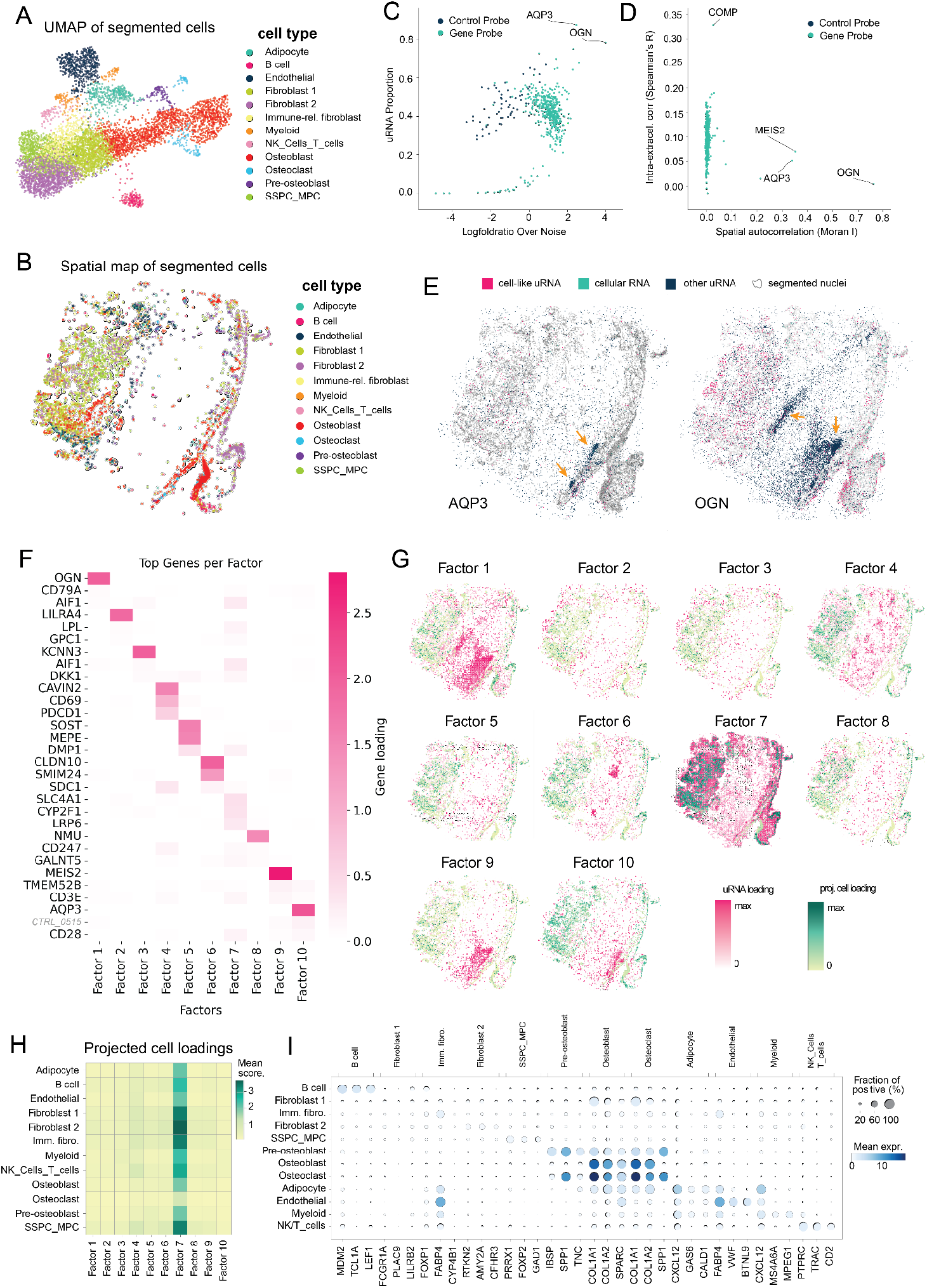
Analysis of the uRNA distribution in the human femur identifies patterns not captured in segmented cells. **(A)** UMAP of the cells segmented in the Xenium human bone dataset, using only nuclear masks. Cells are colored by cell type. **(B)** Spatial map of the cells segmented in the Xenium human bone dataset (panel A), colored by cell type. **(C)** Scatter plot representing, for each type of probe, its uRNA abundance (x-axis), represented as logfold ratio over noise threshold, and its uRNA proportion (y-axis). Dots are colored depending on whether they are control probes or not. **(D)** Scatter plot representing, for each type of probe in the uRNA pool, its spatial autocorrelation (x-axis) compared to its cell-body-uRNA expression correlation for each gene, calculated as the Spearman correlation (R) between expression inside segmented cells and in the surrounding unsegmented space (y-axis). Dots are colored depending on whether they are control probes or not. **(E)** Spatial maps of the Xenium human bone sample displaying transcripts corresponding to AQP3 (left) and OGN (right). Transcripts are colored by RNA status: cellular RNA, or uRNA. uRNA is further categorized based on spatial context into cell-like uRNA (located in regions resembling cells) and other uRNA. Cell boundaries from Xenium’s nuclei segmentation are shown in the background for reference. Yellow arrows are included in each map to highlight accumulations of each type of transcript. **(F)** Heatmap showing the gene loading for all factors of the top three genes with the highest loadings for each factor, obtained from NMF-analysis performed on precomputed uRNA bins in the Xenium human bone dataset. **(G)** Spatial maps showing the factor scores for each uRNA bin across all NMF-derived factors detailed in F (pink scale). Segmented cells are overlaid and colored according to their corresponding normalized projected factor scores (green scale). **(H)** Heatmap representing the mean projected cell loadings of each NMF factor on each of the cell types obtained from the cell type annotation of segmented cells. **(I)** Dot plot illustrating the expression of the top 3 differentially expressed genes for each cell type annotated in segmented cells, across cell types. Dot size represents the fraction of positive cells per gene, whereas the color indicates the mean expression of the gene in each cell type.

**Extended Data Figure 7.**
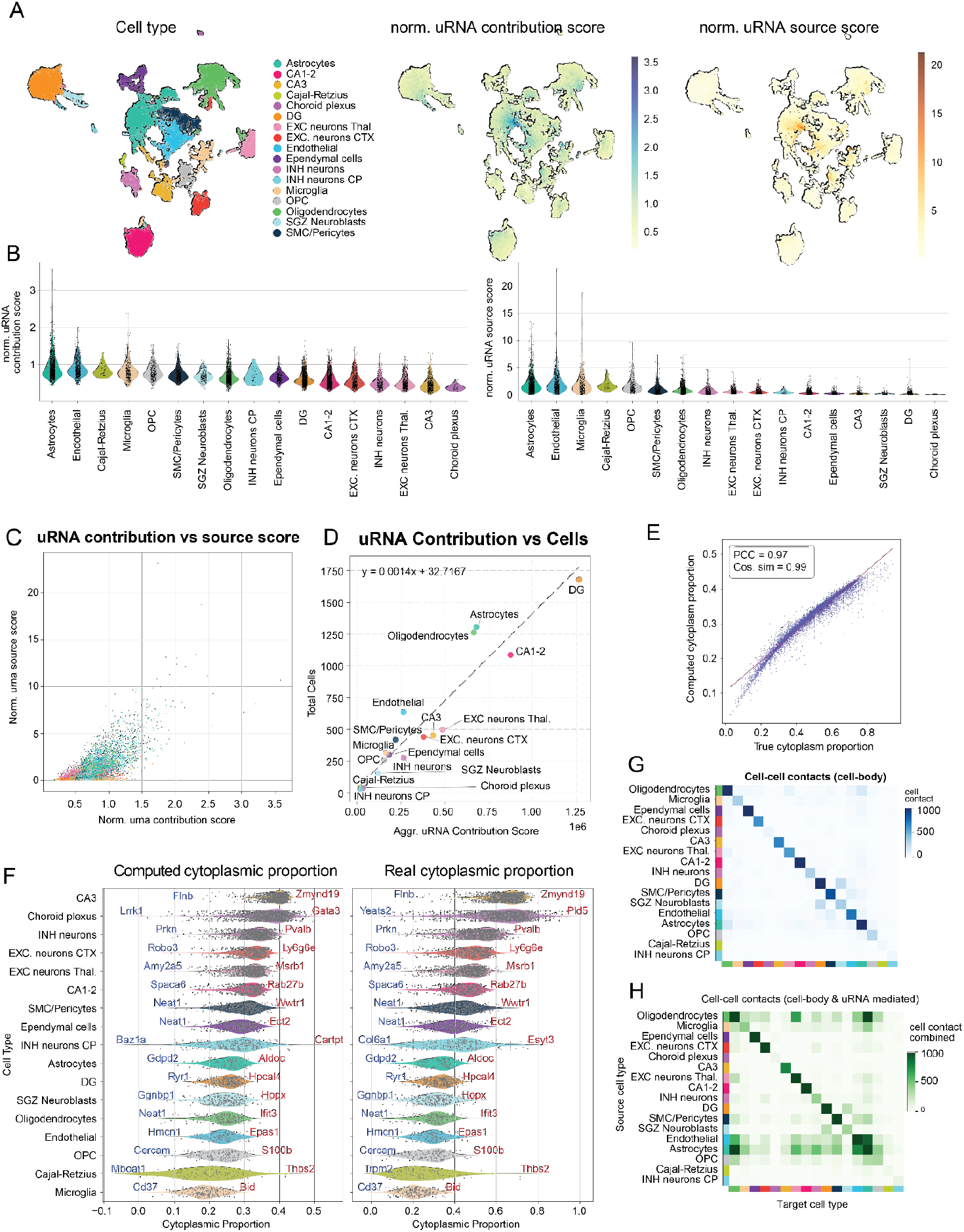
Extended analysis on cell-centric metrics to understand uRNA significance. **(A)** UMAP representations of the cells segmented in the Xenium mouse brain dataset, with cells colored by cell type (left), normalized uRNA contribution score (mid) and normalized uRNA source score. **(B)** Violin plots representing the distribution of uRNA cell-centric scores by cell type, including normalized uRNA contribution score (right) and normalized uRfNA source score (left). A strip plot with the score of each cell, grouped by cell type is overlaid on of the violin plots. **(C)** Scatter plot representing, for each segmented cell in the Xenium mouse brain 5k dataset, its normalized uRNA source score (y-axis) and its uRNA normalized uRNA contribution score. Cells are colored by cell type, as in A. **(D)** Scatter plot representing, for each cell type, the total number of cells detected in the Xenium mouse brain 5k dataset (y-axis) compared to the aggregated uRNA contribution score per cell type. A fitted regression line is included in the plot. **(E)** Scatter plot representing, for each gene in each cell-type, its true cytoplasmic proportion (x-axis) compared to its predicted one (y-axis). **(F)** Violin plot representing, for the genes expressed on each cell type, their real (right) and computed (left) cytoplasmic proportion. Individual dots represent individual genes. The name of the most cytoplasmic and the most nuclear gene in each cell type are overlaid. **(G)** Heatmap showing the number of cell-cell contacts between cell types detected in the Xenium mouse brain dataset using only cell-body positions. **(H)** Heatmap showing the number of cell-cell contacts in total identified between different cell types combining both cell body-based contacts and additional cell-contacts identified using unassigned RNA.

